# Why do we seek information about the future? On the origins of subjective value without instrumental value

**DOI:** 10.64898/2026.06.09.731206

**Authors:** Ethan S. Bromberg-Martin, Josh Merel, Ilya E. Monosov

## Abstract

Why do we want to know the future? Humans and many animals pay for information to predict uncertain rewards, even when they cannot control them. The reason for this conserved yet seemingly paradoxical preference – “subjective value without instrumental value” – remains unknown. Here we develop a normative framework to explain the origins of such preferences, their persistence across species, and their contributions to survival. We formalize and evaluate theories that explain subjective value as originating from an adaptive estimate of advantage for solving core computational problems in naturalistic environments. We show that human and animal subjective values are remarkably well suited to solve these problems, accomplishing the goals of multiple theories simultaneously. We derive novel forms of information to enable existing theories to be dissociated, and show that pooling their subjective values improves performance across diverse environments. Thus, organisms may value information because it pays diverse dividends in nature.

## INTRODUCTION

Humans and many animals often seek *non-instrumental information* about future rewards: information about the future that they cannot use in any way to make better choices or get better outcomes^1^. This appears conserved across many species, including humans, monkeys, rats, mice, and pigeons^2–7^. The price they pay for information, a quantity called the *subjective value of information* (SVOI)^8^, can be remarkably high, sometimes a large fraction of the total available reward, and persists in the face of extensive experience and even explicit instructions that the information is non-instrumental^7–11^. Thus, these organisms assign this information “subjective value without instrumental value.” This form of information seeking has been considered a key signature of curiosity and its evolution across species^12^. It been intensely researched especially in recent years, producing numerous models proposing detailed psychological and neural processes that could generate this behavior. This includes carefully crafted forms of response conflict^13^, conditioned reinforcement^14–16^, uncertainty resolution^5,17–21^, selective attention or salience^22,23^, task engagement^24^, information gaps^25^, frustration^26,27^, anticipatory emotions^28^ including savoring^29^, anxiety^30^, hope^31^, and other affective results of belief states^32,33^, and mixtures of multiple processes^34–36^.

However, little is known about two crucial, fundamental questions underlying this behavior. Why does the brain have these information seeking processes in the first place? And what adaptive functions, if any, did information seeking originate to perform^6,37,38^? These questions are outside the scope of most existing frameworks for modeling psychological and neural processes because they aim to model a process’ inner workings, not the reason why it originated. These questions are also hard to answer for most existing frameworks that attempt to explain why behaviors occur based on decision theory and reinforcement learning, because they aim to use optimality principles to derive behaviors that would improve or maximize instrumental value (how an organism can collect the most rewards from its environment^39,40^), which cannot explain non-instrumental preferences. Furthermore, information seeking is not alone in this regard. It belongs to a large class of non-instrumental preferences, including for information about the future^1^, predictability, controllability, and agency^41,42^, tastes in the absence of nutrients^43^, specific outcome probabilities and sequences^44,45^, viewing novel objects^46–48^ above and beyond their instrumental value for learning and decisions^40,49,50^, and more. Hence, the challenge we face in explaining “subjective value without instrumental value” is not limited to information seeking alone. To address this, we need a general framework to understand the origin of subjective value (above and beyond instrumental value), why subjective values persist across human populations and multiple animal species, and why they may vary across species and individuals.

Here we develop a normative framework to address these questions. We formulate and evaluate theories that explain “subjective value without instrumental value” as originating from adaptive strategies tuned to natural environments. These adaptive strategies could arise from several sources, including evolutionary processes over generations and developmental learning within a single organism’s lifetime. This key logic we formalize here is a well-known phenomenon in evolutionary biology: evolution tends to favor individuals that assign subjective value to options in a manner that approximates the true instrumental value that similar options have in natural environments^6,12,37,38,51^. These species then continue to use those subjective value computations in modern day environments and laboratory experiments, even when the same factors may no longer be accurate cues to instrumental value. For example, many species whose natural diets are high in simple sugars have a sweet tooth, to the degree that they will even pay for artificial sweeteners that lack nutrients; conversely, species like cats or colobine monkeys whose natural diets are low in simple sugars are often insensitive to sweet taste^52–56^. A similar line of reasoning applies to development within an individual organism’s lifetime. For example, humans can learn coping behaviors that are functional to handle stress in early life but persist in adulthood despite becoming maladaptive^57^. Thus, subjective values measured in experiments can be a vital clue to instrumental values in the environments where species evolved and individuals developed (Fig S1).

We then apply our framework to investigate information seeking. We first compile leading extant theories for the origin of information seeking. Each theory proposes that information helps organisms solve a specific computational problem posed by natural environments: the *Time Allocation* problem in foraging (which we refer to as TA theory^37^) or the *Credit Assignment* problem in reinforcement learning (CA theory^6^). We implement each theory by formulating a computational model where an optimal Bayesian agent uses information instrumentally to maximize its reward rate in a naturalistic environment. We then derive the factors that govern the instrumental value of information in each model (IVOImodel), and test whether they predict the factors that govern SVOI in real organisms measured in experiments across species (SVOIexp) using a multi-attribute economic choice task^8^.

Using this approach, we find evidence that humans and animals compute the subjective value of information in a manner that aligns remarkably well with its adaptive value in naturalistic environments. Furthermore, these value computations are surprisingly robust, in the sense that they are well suited to solve multiple computational problems simultaneously in a range of diverse environments. We then leverage our framework to demonstrate that existing theories for the origin of information seeking can be dissociated from each other, by deriving novel types of information for which the theories make divergent predictions. As a consequence, we show that subjective values based on any single theory are the most adaptive in only a narrow range of environments, while mixed value computations incorporating aspects of both theories achieve high performance across diverse naturalistic environments. Thus, our work provides new evidence that organisms compute the subjective value of information in a manner that is non-instrumental in experimental tasks but is conserved across species and well suited to guide adaptive behavior across a range of natural environments.

## RESULTS

### Non-instrumental information seeking is valued and conserved

We develop our framework for investigating the origins of subjective value by focusing on the specific case of *seeking information about the size of an uncertain future reward*. We and others have measured the behavioral preference for this information with a variety of experimental designs, generally with versions of a well-established non-instrumental information seeking task (Fig 1A)^1,3,27^. Participants are offered a choice between two options, Info and Noinfo. If they choose Info, they are shown an informative cue that indicates the size of the upcoming reward *T_advance_* seconds in advance of its delivery (red). If they choose Noinfo, they are shown a non-informative cue leaving them uncertain about the reward size (blue). Importantly, any preference for Info in this task is non-instrumental because there is no way to use the information to control the reward or any other events in the task. Both Info and Noinfo options give a reward *r* drawn from the same distribution *p*(*r*) at the same time *t_outcome_*.

**Figure 1.**
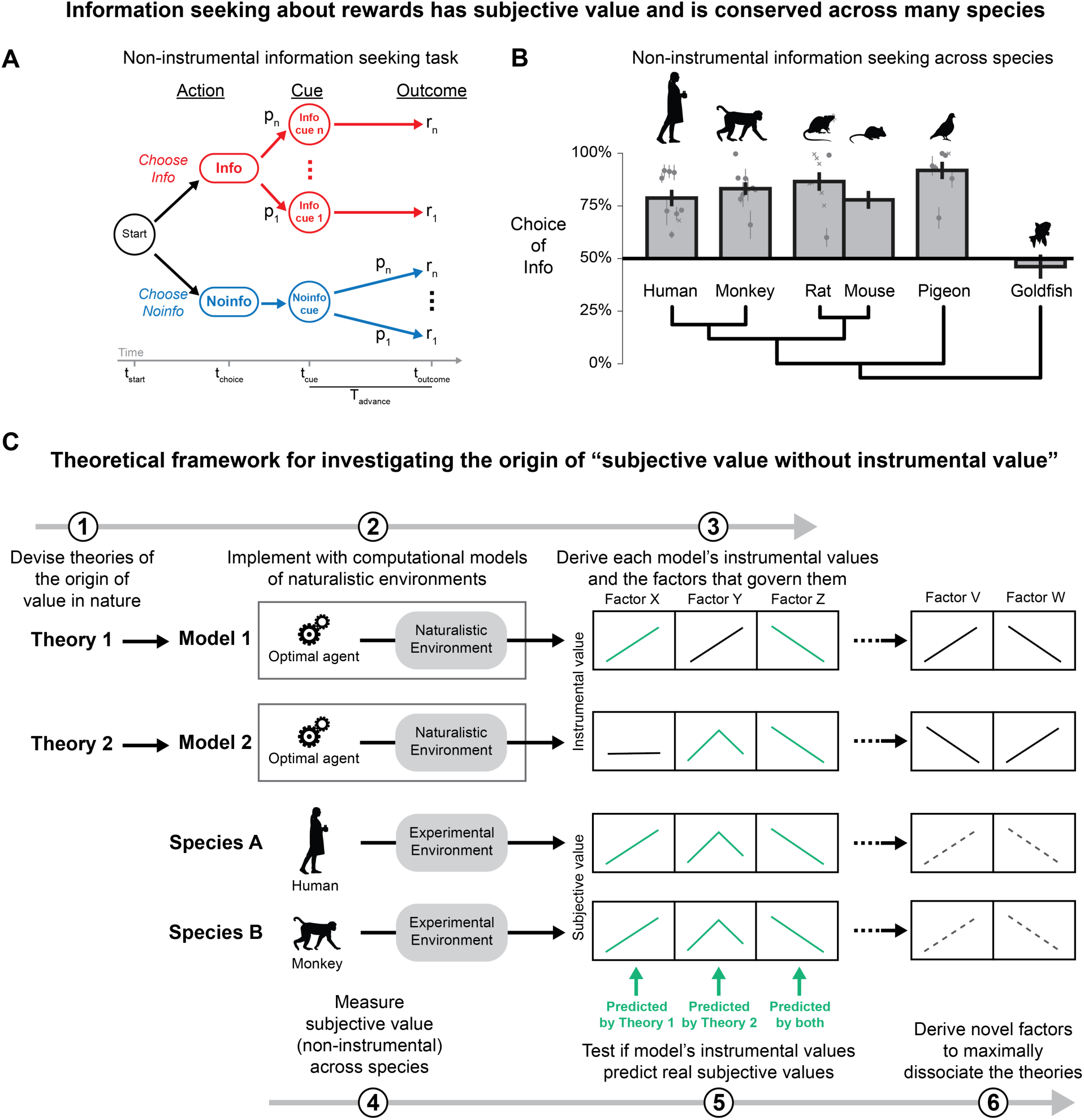
Theoretical framework for investigating the origins of “subjective value without instrumental value,” including information seeking. **(A)** Non-instrumental information seeking task. **(B)** Conservation of information seeking across species. Choice of Info is plotted from a set of 36 datasets across 6 species (Table S1). Bars indicate mean ±SE across data points. Data points are circles with error bars (SE) or crosses (studies not reporting SE). Monkey data is per-animal; mouse and goldfish data are single reports^7,91^. Phylogenetic tree follows NCBI taxonomy (phyloT v2)^107^. **(C)** Theoretical framework. We (1) devise theories for the origin of subjective value in nature, (2) implement each theory with a computational model of an optimal agent in naturalistic environments, (3) derive each model’s instrumental values and how they vary with a suite of motivational factors, (4) measure non-instrumental subjective values across species, (5) evaluate the theories by testing if the model instrumental values predict real subjective values, (6) derive novel motivational factors to better dissociate the theories.

A robust Info preference has been documented in humans and several other species including monkeys, rats, mice, and pigeons, while this preference has been reported to be absent in a more taxonomically distant species, goldfish (Fig 1B). Many individuals in these species place subjective value on information (SVOI) measured by their willingness to pay^7–11^. This occurs consistently across diverse task designs, stimulus and response formats, and rewards^1,27^ (including food^2^, water^58^, money^4^, relief from humid heat^59^, viewing attractive pictures^29,60,61^, etc.).

### Framework for investigating theories of subjective value without instrumental value

To shed light on why this information seeking behavior occurs, we propose a framework for constructing theories of “subjective value without instrumental value” and evaluating their ability to explain behavior (Fig 1C), by linking theory, computational modeling, and state of the art experiments measuring subjective values and the factors that govern them across species. The key logic is that adaptive strategies tuned to natural environments should tend to compute the subjective values of options in a manner that approximates the instrumental values that similar options provide in nature^6,12,37,51^.

We first compile theories for the origin of subjective value (**step 1**) and formalize each theory using a computational model representing its proposal about how using these values to guide behavior helps to solve problems posed by natural environments (**step 2**). We implement each model in the same basic format: an optimal Bayesian agent acts to maximize its reward rate in a naturalistic environment. This allows us to measure the instrumental value of information in each model (IVOImodel) in a simple, direct manner: the improvement in the agent’s reward rate from gaining access to information. We use this to derive quantitative predictions (**step 3**) about the instrumental value organisms can gain in naturalistic environments, and about the key motivational factors that govern this instrumental value (Fig 1C). In parallel, we measure subjective value of non-instrumental information experimentally (SVOIexp) across species, including humans and animals, using standard methods from psychophysics (**step 4**) and determine the key motivational factors these species use to compute subjective value (**step 5**).

With these results in hand we can then evaluate theories against data, by asking which theorized benefits in nature predict actual preferences in the lab. Specifically, we evaluate how each theory’s IVOImodel predicts the observed SVOIexp and how it varies across species and motivational factors (Fig 1C). This comparison can produce several patterns of results with different implications for the origin of subjective value. All factors governing subjective value may be accurately predicted by a single theory; or different factors may be predicted by different theories, suggesting they may originate from different sources. A single factor may be predicted by multiple theories, suggesting that behavior is well suited for multiple purposes; or a factor may not by predicted by any existing theory, suggesting new theories are needed. Importantly, our framework also enables us to use the models to derive novel motivational factors in order to more strongly dissociate existing theories (**step 6**). Thus, as we evaluate theories for the origin of subjective value, we can also use them as guides to identify novel factors that govern value and generate new testable predictions.

In this study we will measure SVOIexp using data from our recent work in humans and macaque monkeys^8^, two closely related primate species. This dataset has three critical features for testing theories of the origin of information seeking. First, this dataset measures SVOIexp across species using closely analogous tasks. This is crucial to test if subjective value judgements are conserved (Fig 1C, green) by reducing the chance that apparent differences in species actually result from differences in tasks. Second, this dataset goes beyond simply estimating a single measure of value per species, by further measuring how individuals in each species scale SVOIexp based on a broad suite of motivational factors that commonly vary in natural environments (including expected reward, multiple forms of reward uncertainty, and information and reward timing). This is crucial to test if theories robustly predict subjective values across varying environments (Fig 1C). Finally, this dataset measures preferences with subjective values rather than more typical choice percentages. This is crucial because subjective values are a more robust measure of preference (Fig S2), and theories of value make their predictions in terms of value-based economic tradeoffs (e.g. how much reward each piece of information is worth^37,62^).

To begin, we first explain the intuitions behind extant theories that have been proposed to explain the origin of information seeking, and implement each of these theories in our framework as a computational model of an agent maximizing its reward rate in a naturalistic environment (Fig 2). Each theory proposes that information’s subjective value originates from its benefit for solving a specific, ubiquitous computational problem posed by natural environments, that presents a core challenge for both biological organisms and artificial agents to overcome.

**Figure 2.**
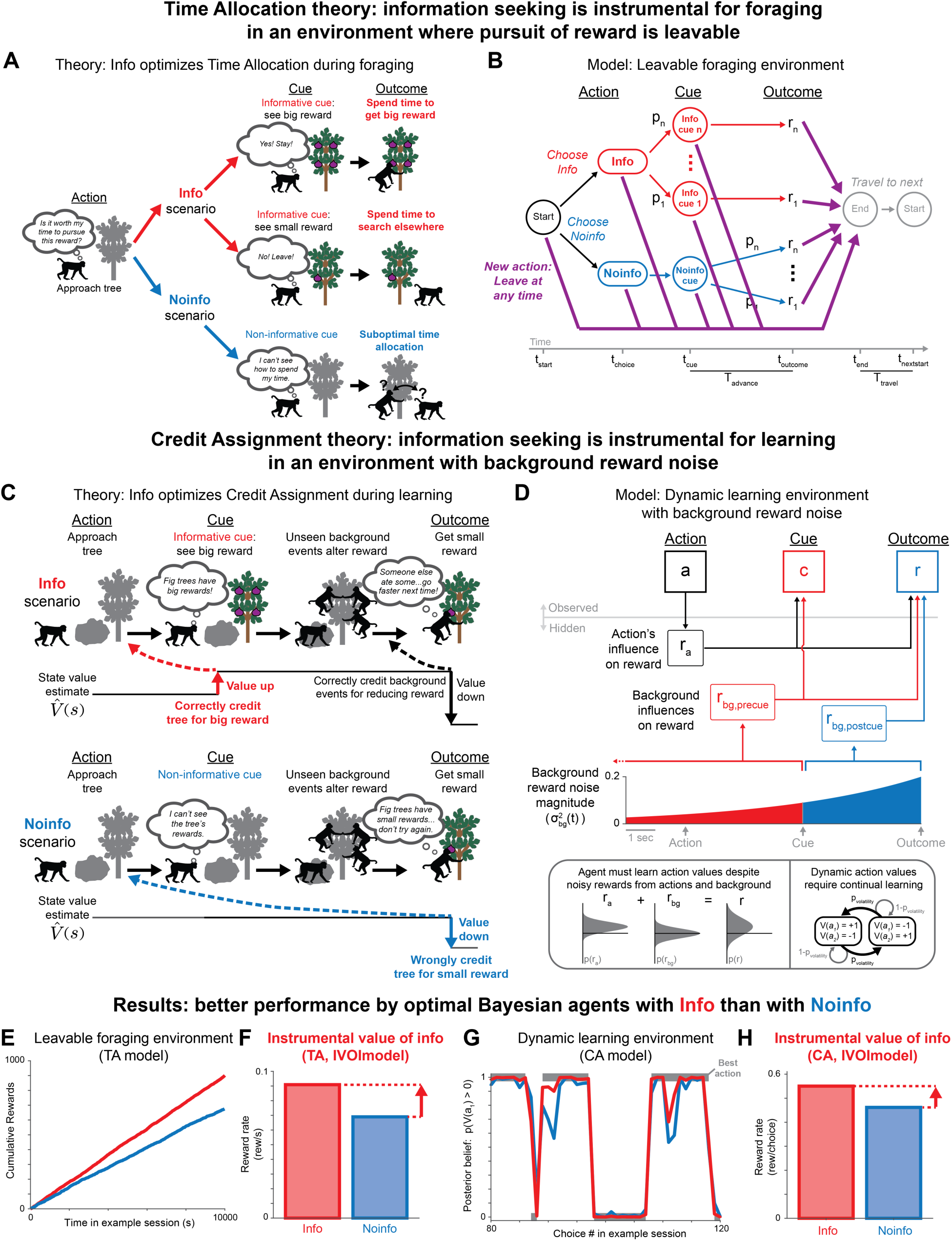
Time Allocation (TA) and Credit Assignment (CA) theories for the origin of information seeking. **(A)** TA theory. Organisms pursuing rewards use Info to decide to *stay* and collect rewards (top, “Spend time to get big reward”) or *leave* to skip them (middle, “Spend time to search elsewhere”), thus getting higher reward rate than with Noinfo (bottom, “Suboptimal time allocation”). **(B)** TA model. An optimal foraging agent pursues reward in a leavable foraging environment. Each state has an extra action to leave the pursuit (purple), which transitions into traveling toward the next pursuit (at a cost of T_travel_). **(C)** CA theory. Organisms use Info to credit actions for their consequences. *Top*: monkey with Info correctly credits the action for a high value state (“Value up,” red), then correctly credits further changes in state value to later background events, even though they are unobserved (“Value down,” black). *Bottom*: monkey with Noinfo cannot observe how the action changed state value, and wrongly credits the action for producing a small reward (“Value down,” blue). **(D)** CA model. Dynamic learning environment with background noise. *Top:* An agent takes an action, gets a cue, then gets a reward. The total reward *r* is the sum of reward from the action *r_a_* plus noise from background events that unfold over time *r_bg_*. The cue indicates state value, including the pre-cue background (red) but not post-cue (blue). *Bottom:* learning is hard due to noisy sources of reward (left) and dynamic action values requiring continual learning (right). **(E)** TA model results. Cumulative reward over time in an example simulated session is higher for an **o**ptimal agent with Info (red) than Noinfo (blue). **(F)** Instrumental value of info (IVOI, red arrow) for the TA model is the difference in reward rate between Info vs. Noinfo (red bar vs. blue bar). **(G)** CA model results. Optimal agent learns better with Info than Noinfo (red vs. blue), shown by the time course of posterior belief that action *a_1_* is good over 40 example choices. The best action during each choice is shown in gray. The agent with Info changes its belief more quickly when the best action changes, and stays closer to that belief while the best action is stable. **(H)** IVOI for the CA model.

### The Time Allocation theory of information seeking

TA theory proposes that information seeking is adaptive behavior to solve a central problem in optimal foraging theory: foraging animals must continually decide whether the pursuit of a reward is worth their time^37^ (Fig 2A). This is because animals in natural environments have the option to abandon their pursuit at any time^63^ (unlike non-instrumental lab tasks where there is no option to leave a trial early; Fig 1A).

This means information about the size of future rewards has high instrumental value (Fig 2A, “Info scenario”)^37^. If the animal gets ‘good news’ that the reward is big, they can decide to *stay* in pursuit, allocating their time to collect the reward. If the animal gets ‘bad news’ that the reward is small, they may decide to *leave* the pursuit, allocating their time to search for better opportunities. If the animal gets no information, they will allocate their time suboptimally because they will not know when to stay or leave (Fig 2A, “Noinfo scenario”). Importantly, this prediction of TA theory contrasts with influential classic theories of optimal foraging, which do not consider information seeking because they make the simplifying assumption that animals have perfect knowledge about future rewards in their local environment^62,63^.

To formalize this in a model (Fig 1C, step 2), we implement an optimal Bayesian agent to maximize reward rate in an environment consisting of a Markov decision process representing pursuit of reward (Fig 2B, Methods). Ideally, we would design the model based on the detailed statistics of informative cues and rewards in natural environments; however, little is known about these natural statistics, and methods to measure them are only now being developed^64^. Therefore, we design all models in this paper based on the simplest, most generic possible scenario: an agent takes an action, observes a cue, and gets a reward outcome. This scenario is broadly applicable because it commonly occurs for organisms in natural and modern environments, as well as for artificial agents, and can be directly mapped onto experimental information seeking tasks (Fig 1A) and both TA and CA theories (Fig 2). We start by modeling a standard laboratory task environment where information is non-instrumental (Fig 1A), where agents achieve the same reward rate from Info and Noinfo. We then expand our formulation to a more naturalistic foraging environment where the agent can leave its pursuit at any time and immediately start searching for the next reward to pursue (Fig 2B, “New action: Leave at any time”). The agent now gains a higher reward rate from Info than Noinfo (Fig 2E) resulting in positive IVOImodel (Fig 2F).

The optimal agent indeed gains value from information through the time allocation mechanism proposed by the theory (Fig S3). Specifically, the agent decides to stay or leave by comparing the value of its current state, V(s), against the average reward rate from the global environment as a whole, *RRavg*. In most of our simulated environments we simply set *RRavg* to a fixed value, but we replicate a similar pattern of results if we set *RRavg* adaptively based on the agent’s behavior (Methods). Finally, note that our formulation allows agents to leave at any time from any state (Fig 2B). This extends previous work which considered the special case where agents can only leave upon entering the cue state^37^. This is necessary to model naturalistic environments where agents can abandon pursuits freely, and changes how motivational variables influence IVOI (Fig S4).

### The Credit Assignment theory of information seeking

CA theory proposes that information seeking is adaptive behavior to solve a central problem in reinforcement learning: an agent must decide how much credit its past actions deserve for producing future rewards^6^ (Fig 2C). If an action was responsible for getting more reward, the agent should learn to choose it more; if the action was responsible for getting less reward, the agent should learn to choose it less. This is called the temporal credit assignment problem, and it is a difficult dilemma with no general solution^65,66^. The temporal credit assignment problem is harsh in natural environments because each action is only one of many causal influences on future rewards. Other influences unfold continually over time, including the agent’s other actions, actions by other agents, hidden environmental states, inherent reward variability, and so on. This ambiguity makes it hard to learn action values.

Crucially, CA theory points out that agents can improve credit assignment by disambiguating these potential influences on reward, by using information about future rewards to see how state values change over time^6,66^. To see this, suppose an agent takes an action and immediately sees an informative cue giving ‘good news’ that a big reward is available. The agent can estimate that the expected future reward value of its current state has increased (Fig 2C, “Info scenario”). This triggers a positive reward prediction error (RPE) that the agent correctly credits to its action to learn it has a high value (Fig. 2C, “Value up”). Then, even if later events interfere – like competitors consuming part of the outcome, causing the agent to only receive a small reward – the agent can accurately credit this change in state value to later background events in the environment, not to its original action (Fig 2C, “background events alter reward” → “Value down”). In other words, the agent can infer “My action was good because it brought me to a high value state. I only got a small reward because background events must have interfered.” On the other hand, suppose the same agent sees a non-informative cue. The agent would not know that its action led to a high value state and would only observe the final small reward, so it would wrongly credit its action for the low value outcome (Fig 2C, “Noinfo scenario”). The agent would wrongly infer “My action was bad because it got me a small reward.”

We note that this ability to improve credit assignment using state value information is a foundational principle in reinforcement learning. For example, the influential TD learning algorithm^40^ was designed with this principle as its primary improvement over older methods^66^ leading to advances in machine learning^67^. This principle is also thought to explain why animals evolved to use similar learning algorithms, such as using RPEs as teaching signals^68^ which are implemented by neural systems including the dopaminergic reward pathway that produces reinforcement^69^. The CA theory extends these observations by proposing that organisms use adaptive information seeking behavior to support these learning algorithms, by placing subjective value on information that improves CA in natural environments^6^.

To formalize this in a model (Fig 1C, step 2), we implemented a model with the three necessary parts of the theory: (1) a standard dynamic reinforcement learning environment, (2) where both the agent’s action and background events influence the reward, thus creating a CA problem, (3) which agents can partially solve, using informative cues indicating state values (Fig 2D). Again, because little is known about the statistics of these events in natural environments, we designed the CA model to capture the core intuition of the theory in the simplest, most generic possible scenario. Thus, we model a single action, cue, and reward, and a single “background process” representing all other events that influence the reward. This compact model formulation is also necessary to make it tractable to derive the optimal Bayesian learning agent (since the CA problem lacks a known optimum in the general case^66^).

In the reinforcement learning environment, an agent repeatedly takes an action *a* which gives a reward *r_a_* drawn from a distribution p*_a_*(*r_a_*) (Fig 2D). There are two possible actions, *a_1_* and *a_2_*. One is a *good* action drawing rewards from a distribution with mean = +1. The other is a *bad* action with an equal and opposite distribution with mean = -1. Identifying the good action is difficult because the reward distributions are noisy (Fig 2D, bottom left), and the action values are dynamic, occasionally reversing (Fig 2D, bottom right).

To model the CA problem, we include the naturalistic feature that the reward is also influenced by background events unfolding over time, modeled by adding Gaussian noise to the reward at each moment (Fig 2D, “Background influences on reward”). Thus, the agent gets a total reward that is the sum of the action’s contribution plus the background’s contribution: *r* = *r_a_* + *r_bg_*.

Crucially, the agent only observes the total reward *r*. Thus, the agent must decide how much of *r* to credit to its action vs. the background. The background noise ramps up to the time of reward delivery with a simple exponential function (Fig 2D) to model the classic principle of temporal contiguity in natural environments, that events occurring closer to the outcome tend to have greater casual influence^70^. Our results do not depend on the precise timecourse and hold for a wide range of noise parameters (Fig S5, Methods).

Finally, to model information, we include a cue *c* indicating the reward value of the current state (Fig 2D, “Cue”). Specifically, the cue indicates the reward contributed thus far: the action’s contribution plus the part of the background’s contribution occurring before the cue: *c* = *r_a_* + *r_bg,precue_* (Fig 2D, red). Thus, the agent can partly solve the credit assignment problem by learning from the cue instead of the reward. Essentially, the cue provides a better teaching signal than the reward (Fig S3). The cue has the same ‘signal’ about the action’s value (reward from the action) but less ‘noise’ (reward from the background) because it excludes the portion of the noise that occurs after the cue (*r_bg,postcue_*; Fig 2D, blue shaded area). Indeed, the optimal Bayesian agent has superior performance with informative cues: more stable and accurate beliefs when the environment is stable, and faster belief updating when action values change (Fig 2G). The agent gets a higher reward rate with Info resulting in positive IVOImodel (Fig 2H).

Thus, both TA and CA models produce IVOImodel in naturalistic environments, providing candidate explanations for information’s subjective value. However, they predict that information value arises for very different purposes. Therefore, we next use our framework to evaluate these theories, by deriving their predictions about how a broad suite of motivational factors influence IVOImodel and testing them against data.

### Both theories explain information value scaling with an SD-like form of uncertainty

Humans and animals potently regulate SVOIexp with the features of information and reward^8,9,11^. We first tested if the TA and CA theories explain how SVOIexp scales with two prominent motivational factors: expected reward and reward uncertainty (E[r] and Unc[r]; Fig 3A).

**Figure 3.**
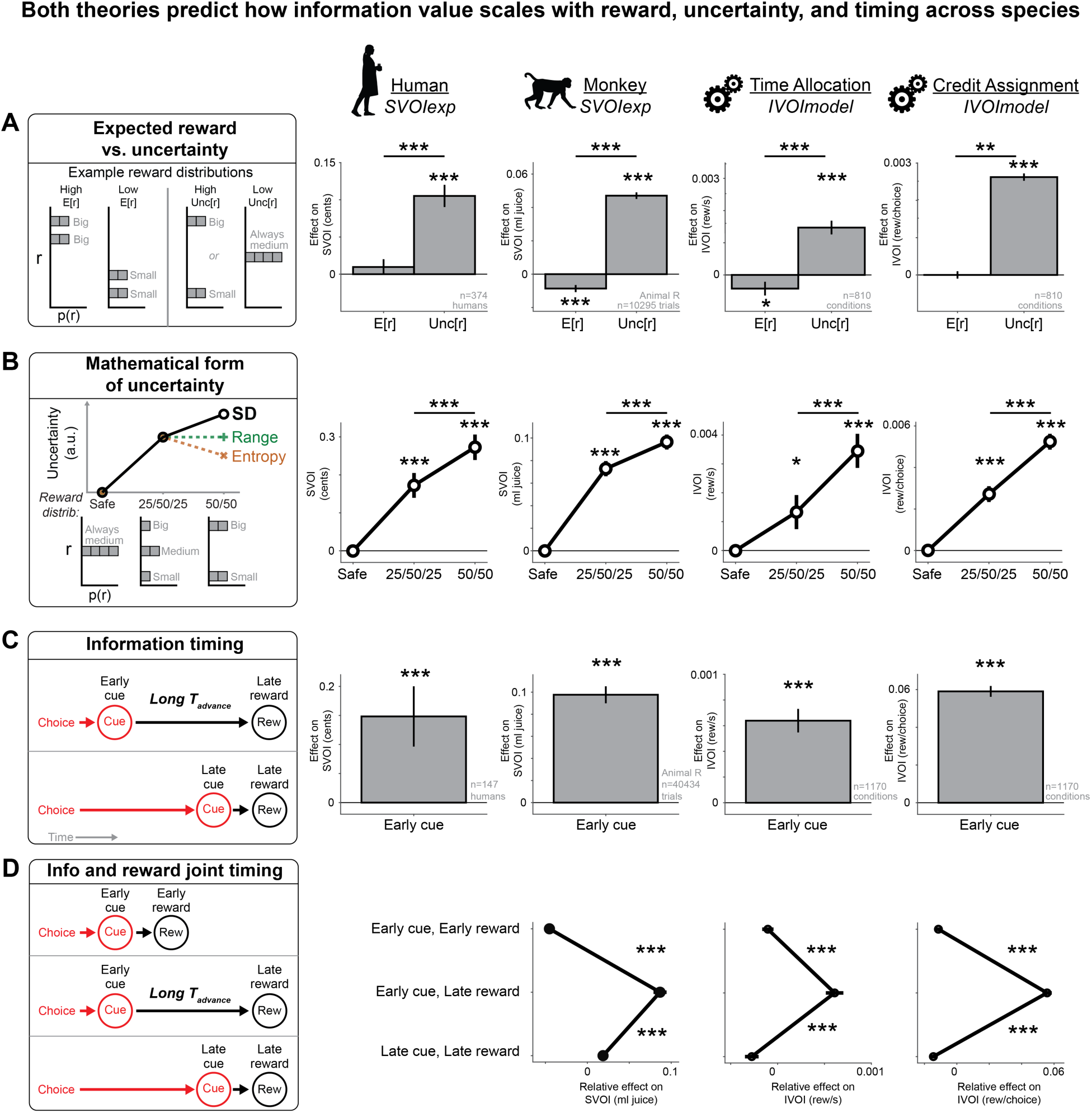
TA and CA theories predict information value scaling with reward, uncertainty, and timing across species. **(A)** Expected reward vs. uncertainty. *Left:* Concept for manipulating E[r] vs Unc[r]. Example reward distributions where reward sizes are: big, small, big or small, or medium. *Right:* Mean effects of standardized E[r] and Unc[r] on SVOIexp in the human population and an example monkey, and IVOImodel from the TA and CA models, expressed in units of reward. Unc[r] effect is larger than E[r]. Error bars are ±SE. *, **, *** indicate p < 0.05, 0.01, 0.001 (humans: signed-rank test of weights, or difference of weights, vs. 0; others: *t*-test of GLM weights, difference of weights, vs. 0) **(B)** Mathematical form of uncertainty. *Left:* uncertainty measures resembling SD[r], Range, or Entropy have higher, equal, or lower uncertainty, when comparing a 50/50 reward distribution to a 25/50/25 distribution. *Right:* Mean effects of 25/50/25 and 50/50 distributions on SVOIexp and IVOImodel. 50/50 distributions have significantly higher effect. **(C)** Information timing. *Left:* information comes earlier in advance (long Tadvance, black arrow) if the cue is early (top) rather than late (bottom), given fixed reward time (Rew). *Right:* Positive mean effect of early cue (vs. late cue) in SVOIexp in the human population and an example monkey, and IVOImodel from the TA and CA models. **(D)** Joint information and reward timing. *Left:* information comes earliest in advance with an early cue and late reward (middle), compared to both early or both late (top or bottom). *Right:* Mean relative effect of timing categories on SVOIexp and IVOImodel. Early cue and late reward has greater value. Error bars are ±SE.

We measured IVOImodel in a range of simulated environments that independently varied E[r] and Unc[r] (Methods). We measured SVOIexp using the choices of n=374 humans and n=4 monkeys performing analogous non-instrumental information seeking tasks that independently varied E[r] and Unc[r] for each offered option^8^ (Methods; Fig S6). Humans worked for money as a reward, while monkeys worked for juice. We defined each individual’s SVOIexp in the standard manner, as the reward they paid for information (by choosing Info offers with lower E[r] than Noinfo offers; Fig S6)^8^. To quantify how IVOImodel and SVOIexp are governed by motivational factors, we fit the subjective value of each offer using standard generalized linear models (GLMs) as a simple linear weighted combination of factors:

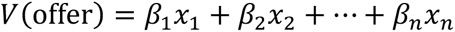

…where *x* is a vector of motivational factors (including E[r], Unc[r], and Info), and interactions (Methods). The key weights in this model to test our hypothesis are the interaction terms representing how the value of information scales with E[r] and Unc[r] (Fig 3A). As in previous work^8^, we defined Unc[r] as the standard deviation of rewards (SD[r]), standardized all scalar regressors, and re-scaled the weights to be in units of reward (Methods).

We find both TA and CA theories make two clear predictions: information value increases with Unc[r], and increases more with Unc[r] than E[r] (Fig 3A, IVOImodel). These results are robust for a wide range of parameters (Fig S5, Methods). Crucially, both predictions hold true for subjective values of humans and monkeys (Fig 3A, SVOIexp; true for all animals, Fig S7).

Furthermore, we find that both theories correctly predict the specific family of mathematical uncertainty measures that humans and monkeys use to value information^8^ (Fig 3B). Most proposals for how organisms measure uncertainty can be grouped into three broad families: functions of reward probabilities (like Shannon entropy^71^), reward sizes (like range^72^), or both probabilities and sizes (like SD or variance^73^). To dissociate these families, we used three reward distribution types: *safe* offers that always give medium reward; 25/50/25 offers that give 25% chance of big, 50% chance of medium, and 25% chance of small, and 50/50 offers that give 50% chance of big or small. Representative measures from each family make distinct predictions about which distribution is the most uncertain^8^ (Fig 3B, left). We find that both theories correctly predict that information value increases with an uncertainty measure more closely resembling SD than the alternatives, with IVOImodel and SVOIexp significantly highest for 50/50 reward distributions for both theories and both species (Fig 3B, right; true for all animals, Fig S7).

Crucially, while TA and CA theories make remarkably similar predictions, they do so for distinct reasons (Fig S3). The TA model’s IVOI strictly grows with Unc[r] because it increases the benefit of correct stay/leave decisions: correct ‘stays’ get bigger rewards while correct ‘leaves’ avoid smaller rewards. By contrast, the CA model’s IVOI is maximized by relatively high Unc[r] because it creates a high baseline noise level so that learning is hard without information but becomes easy with information. The two theories also have different reasons for the relatively smaller effect of E[r]. The TA model’s IVOI is maximized when the average value of the pursuit (E[r]) is equal to the average value of the environment (*RRavg*), because then the agent needs information about the pursuit’s specific outcome to decide to stay or leave. As a result, variations in E[r] across offers can have either positive, negative, or zero effect on IVOImodel, depending on whether the average offer value is lower, higher, or equal to *RRavg* (Fig S5). This provides a potential explanation for the substantial variation across individuals and studies in the effect of E[r] on information seeking^1,8,32,34^, which may arise due variations in actual or perceived *RRavg* (Fig 3A, animal R; Fig S7). By contrast, in the CA model E[r] produces an additive offset to reward size that does not affect learning. Thus, while TA and CA theories make similar predictions about Unc[r] and E[r] overall, our models suggest they may diverge in regimes not yet tested experimentally (Fig S5)

### Both theories explain information value scaling with information and reward timing

We next tested if TA and CA theories can explain how SVOI scales with the timing of events, including both cues and rewards (Tcue and Trew). To test this, we measured IVOImodel for each model in a range of simulated environments that varied both Tcue and Trew (Methods). We also measured SVOIexp using the choices of n=147 humans and n=3 monkeys performing analogous non-instrumental information seeking tasks where offers presented cues with variable timing^8^. The human task varied Tcue and the monkey task varied both Tcue and Trew. As before, we estimated IVOImodel and SVOIexp using analogous GLMs, measuring the effects of cue timing (Fig 3C) and joint cue and reward timing (Fig 3D).

We find that both theories make a clear prediction: information value grows when Tcue is early and Trew is late, in other words, when the cue comes long in advance of the reward (Tadvance, Fig 3C,D). Thus, in our test of cue timing, IVOImodel in both models is greater when Tcue is early (Fig 3C). Crucially, the same is true for SVOIexp in humans and every monkey (Fig 3C, Fig S7). In our test of joint cue and reward timing, IVOImodel in both models is greater when Tcue is early and Trew is late, than when both are early or both are late (Fig 3D). Again, the same is true for each monkey (Fig 3D, Fig S7). There is evidence this is true for humans as well. This cannot be tested using our human dataset because participants did the task online, so they could not be constrained to ‘consume’ their monetary rewards at a specified time. However, multiple human experiments reported that non-instrumental information seeking grows when Tcue is fixed and Trew is lengthened, using rewards that are only ‘consumable’ at specified times (viewing attractive pictures^29,60^).

The TA and CA theories again make their similar predictions for distinct reasons (Fig S3). In TA theory longer Tadvance saves the agent time, by letting the agent make its stay/leave decision sooner in advance of the reward. In CA theory longer Tadvance makes the cue a better teaching signal, by making more of the background noise fall into the post-cue period.

### Theories value information distinctly for immediate TA vs. longer-term CA

Our findings thus far indicate human and monkey subjective values may be simultaneously beneficial for solving both TA and CA problems in naturalistic environments. Therefore, we next used our framework to investigate whether these theories make any dissociable predictions. To do this, we turned from *full* information (cues indicating the exact future reward) to the less studied case of *partial* information^74^ (e.g. cues indicating specific features of reward). We derived the minimal necessary partial information for each model to obtain its full IVOI, then compared their reward rates using full vs. partial information in analogous environments (Methods, Fig. 4A,B), to generate experimentally testable predictions about the relative value of partial information (Fig 4C). We find that each model only extracts instrumental value from a small part of the full reward information. Further, they use largely distinct parts of the information. Thus, each model gets maximal IVOI using its preferred information type, and much lower IVOI using the other model’s preferred type (Fig 4C).

**Figure 4.**
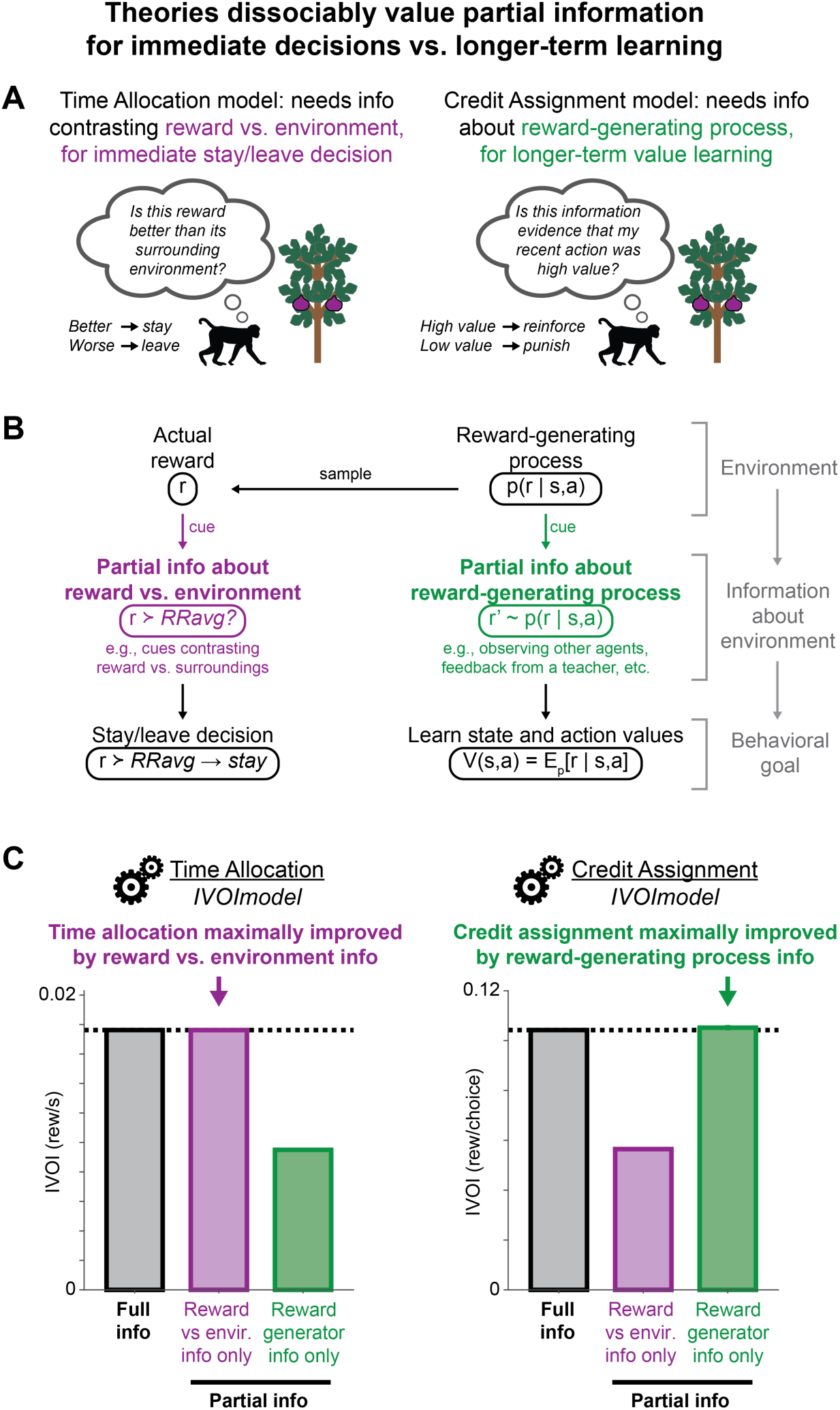
TA and CA theories value dissociable parts of information for immediate stay-leave decisions vs. longer-term learning. **(A)** Theories value dissociable parts of information. *Left:* TA model uses the part of info contrasting the reward vs. its environment for stay-leave decisions. *Right:* CA model uses the part of info about the underlying reward-generating process for value learning. **(B)** Partial information is a binary indicator comparing reward vs. *RRavg* for stay-leave decisions (purple), or a fresh random draw from the reward-generating process for value learning (green). **(C)** Theory predictions: IVOImodel for matched TA and CA models (left, right) given standard full information (black) or partial information types (purple, green). Error bars are ±SE.

The intuition is simple. The TA model only uses information about future reward to aid its immediate decision to stay or leave. Therefore, it does not need to know the reward’s exact features (probability, magnitude, timing, etc.). All it needs to know from the cue is whether the reward is worth pursuing – whether pursuit is better than alternatives in its environment (Fig 4A). As an experimentally testable example, the model can maximize IVOI using a partially informative cue that is a simple binary indicator, comparing the expected value of staying for *r* vs. leaving to obtain *RRavg* (Methods, Fig 4B,C). In nature, organisms may obtain similar partial information from cues contrasting potential rewards vs. their surrounding environments. For example, a visual cue indicating the relative health of the nearest tree vs. surrounding trees reveals whether its fruit are worth pursuing, but not its exact bounty.

By contrast, the CA model only uses information to learn action values to aid its future decisions in the longer term. Therefore, it does not need to know the actual, noisy outcomes of its actions. All it needs from the cue is evidence about the underlying reward-generating process: the true state and action values (Fig 4A). As an experimentally testable example, the model can maximize IVOI using a partially informative cue based on hypothetical, fictive outcomes^75–78^ generated as fresh random draws *r’* from the true outcome distribution (Methods, Fig 4B,C). This information is statistically independent of the agent’s actual reward *r*, but is equally valid evidence to learn the underlying action value. In nature, agents may obtain similar partial information by observing other agents’ outcomes^79^, a teacher’s instructions^80^, or mental simulation of alternatives^81^.

Thus, TA and CA theories make experimentally dissociable predictions about how organisms should value different types of partial information, based on its benefits for solving the TA and CA problems in natural environments.

### Mixed value computations based on both theories succeed in a wide range of environments

The fact that different information types have different uses for solving the TA and CA problems has important implications for the origin of subjective value. Organisms in nature must solve *both* problems, so they should benefit from both information types. Further, organisms rarely have perfect knowledge about their environments, which are diverse and dynamic over multiple timescales, so organisms can rarely compute the precise benefit of solving one problem versus the other.

Therefore, we reasoned that it may not be productive for organisms to rigidly compute SVOI for optimizing TA or CA alone. Instead, they might construct mixed value computations with elements of multiple theories, to handle diverse environments (Fig 5A). For example, suppose an organism computes SVOI using a 70/30 mixture of TA/CA theories. It would perform best in environments where information is mostly useful for TA, but may still perform passably if information is useful for both TA and CA (Fig 5A, environments 1-3). Furthermore, if information seeking is adaptive, then this could result in such mixed value computations becoming strongly adapted to a specific mixture, or else becoming weakly adapted resulting in heterogeneity across species and individuals, depending on how permissive natural environments are toward diverse information seeking strategies.

**Figure 5.**
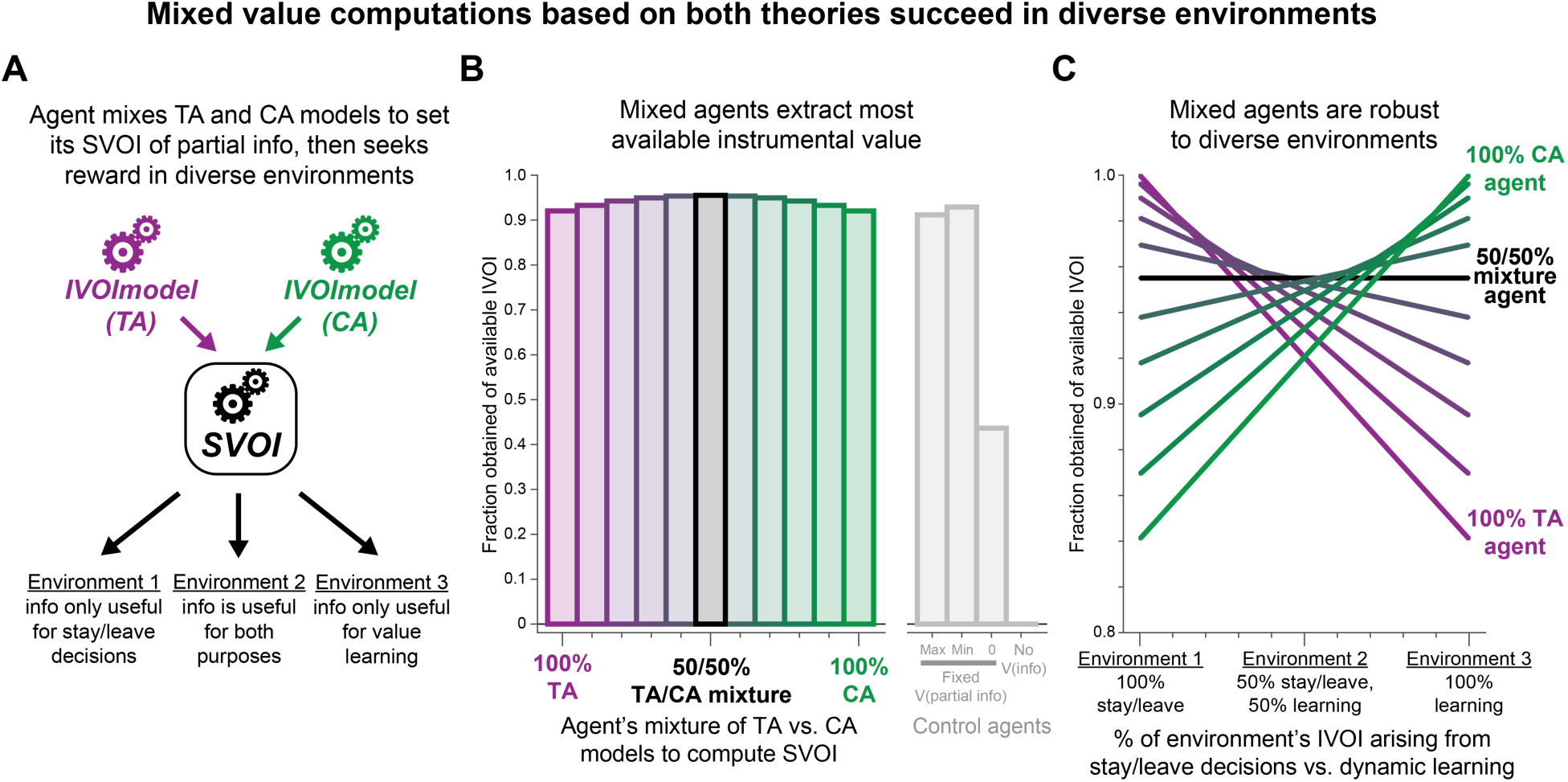
Mixed value computations based on both theories succeed in diverse environments. **(A)** Concept: agent computes SVOI of partial information as a mixture of IVOImodel from TA vs. CA models, then pays for information in environments where different portions of IVOI derive from solving TA vs. CA problems. **(B)** Agents harvest most available IVOI in a set of environments (y-axis) by setting SVOI with mixtures ranging from 100% TA to 100% CA (x-axis, purple to green colors), with maximal performance at 50/50% mixture (black). They generally outperform control agents with the same fixed SVOI for all partial information types (gray). **(C)** Same as B, but each colored bar is now a curve showing performance separately for each environment (x-axis, % environment’s total IVOI from solving TA vs. CA problems; n=11 environments, first/middle/last are environments 1/2/3 from panel A). The 50/50% mixture agent (black) performs well in all environments, while extreme mixtures perform better in their specialties but worse in most environments (100% TA and CA, purple and green).

To investigate this, we model the behavior of a set of agents that compute their SVOI of partial information based on different mixtures of TA vs. CA theories, spanning the full range from 100% TA to 100% CA (Fig 5B,C). Importantly, unlike the agents we examined thus far, these agents are not optimized for any one specific environment. Instead, their SVOIs are based on a simple linear mixture of the theory-derived IVOIs (Fig 4C). We then evaluate each agent’s performance in a diverse set of environments. All environments have the same total IVOI, but have different proportions of it arise from solving the TA vs. CA problems (Fig 5A, environments 1-3). All environments offer agents opportunities to pay for each information type (Fig 4C). We quantify each agent’s performance by its achieved IVOI as a fraction of the environment’s total available IVOI (Fig 5B,C). Thus, an agent achieves optimal performance only when its SVOIs are matched to the true IVOIs of each information type in its environment.

We find mixed value computations are broadly successful in a range of environments. On average across this full set of environments, agents with all possible mixtures extract a large fraction of the maximum available IVOI (Fig 5B, all > 90%). Importantly, these mixed agents do not simply treat all information types the same. They gain their IVOI by adaptively valuing the different information types. This is shown by the fact that almost all these mixed agents outperform control agents that assign identical SVOIs to all partial information types (Fig 5B, gray; all partial information types set to the same maximum, or the same minimum, or to zero SVOI; or all information set to zero SVOI). Overall information seeking performance across all environments was only modestly sensitive to the precise mixture level (Fig 5B).

The most successful agents are intermediate mixtures, which appear to be a ‘sweet spot’ for success in diverse environments. When evaluating agents separately in each environment, the 50% mixture achieves high IVOI in all environments (Fig 5C, black) thus producing the best average performance (Fig 5B, black). More specialized agents extract modestly higher IVOI in extreme environments matching their preference (e.g. 100% TA agent is the best in the 100% TA environment), but at the cost of substantially worse IVOI in balanced and mismatched environments (e.g. 100% TA agent is the worst or tied for worst in the 50% and 0% TA environments) producing worse average performance (Fig 5B, purple).

Thus, we predict that information value computations, and the underlying neuronal circuitry, will not rigidly follow a single theory’s dictates. Instead, they will compute value using a mixture of methods, with potential for considerable diversity across species or individuals that face both TA and CA problems in nature. Our approach will allow this to be tested experimentally by making it possible to infer mixed value computations from choice data (Fig S8).

## DISCUSSION

We introduced a theoretical framework for investigating the origin of subjective value without instrumental value, and applied it to gain insight into a conserved form of motivation: information seeking about uncertain future rewards. We evaluate leading theories for the origin of information seeking that propose organisms value information for solving very different computational problems posed by natural environments: optimizing time allocation for foraging and credit assignment for learning. Yet remarkably, we find that both theories make a series of similar, specific, and correct predictions about information value in both humans and monkeys, suggesting their subjective values are well suited to enhance both time allocation *and* credit assignment. Our results thus provide evidence that subjective value computations across species originated in a manner rendering them surprisingly robust, capable of paying dividends in diverse naturalistic environments.

Our framework formalizes a critical yet understudied avenue of investigation into information seeking: understanding this strikingly conserved behavior in light of its benefits in naturalistic environments^6,37^. Crucially, because this framework explains subjective values as originating from natural environments rather than experimental environments, it provides an intuitive explanation for why humans and animals persistently assign SVOI to non-instrumental information despite extensive experience or explicit task instructions^2,6,8,27,34^, and why their SVOI can be affected surprisingly modestly by changes to information’s instrumental value in experiments^82–85^. This explanation is in distinct from, and complementary to, conventional process models of subjective value. Process models seek to explain the psychological processes and neuronal circuits that compute subjective values, but do not formally model why organisms use those processes in the first place, or what problems they solve in nature. In particular, our framework provides a new avenue to evaluate process models. A process that works well in experimental environments may fall short in nature, and vice versa. Thus, in addition to the conventional method of evaluating process models based on their ability to fit behavior in experimental environments, our framework provides an method to evaluate them in different naturalistic environments, by computing the reward rates they would achieve. Furthermore, we show our framework produces explanations for why humans and animals adjust their SVOIs as they do based on broad suite of motivational factors, including expected reward, a specific mathematical form of reward uncertainty, and cue and reward timing.

In addition, our framework makes it possible to use theories about the origin of subjective values to identify novel motivational factors that neural systems may sense and use to guide decisions. We do this here by deriving the minimal necessary information to compute subjective values that produce maximal performance in naturalistic environments. This provides new insights into the nature of subjective value, and corresponding experimentally testable predictions about the underlying neural computations and behaviors (Fig 4,5). Our results also demonstrate the importance of evaluating theories of value in a common framework. We find that theories about the origin of information seeking are much more closely aligned than was previously known, making many similar predictions. Only by employing our framework – by comparing theories using their IVOImodels - could we identify novel motivational factors that are distinctive signatures of each theory, leading to dissociable predictions.

We also use our framework to show that, if information value originated to serve specific purposes, then organisms may value information sources based on only a small portion of their total informational content. This makes the surprising and experimentally testable prediction that conventional reward cues in experiments may only contain a small kernel of information that organisms value, while the rest is ‘excess’ information. Furthermore, each theory places value on a different portion of the information. TA agents only want to know if it is worth their time to pursue the reward, to make immediate stay-or-leave decisions. By contrast, CA agents only want to learn the underlying state and action values that generated the reward, to improve their decision-making policy in the long term. In short, TA agents want to know “what should be my next action?” while CA agents want to know “what should be my long-term policy?” This makes the experimentally testable prediction that organisms may compute their subjective values based on dissociable types of information. This also suggests that organisms might value information types differently depending on the statistics of their natural environments, and might shift their information seeking strategy adaptively to achieve their current goals (e.g. foraging for scarce food to survive the day vs. learning skills to flourish longer term).

Indeed, our results show that mixed subjective value computations, that incorporate elements of multiple theories, succeed in diverse environments. Subjective values derived from a single theory are optimal in specialized environments matching that theory’s assumptions, but perform substantially worse in environments with diverse information types and uses. This result may explain why SVOI is reported to vary with factors reminiscent of multiple theories. Experiments derived from TA theory suggest pigeons show key signatures of stay-leave decisions, by discounting low value outcomes as though they were leavable^82^, and leaving if an option to leave is available^83^. Meanwhile, experiments manipulating feedback cues suggest that organisms can prefer information that does not directly predict future reward but may aid credit assignment for learning state and action values, such as getting an early peek at half of a configural cue^86,87^ or information about counterfactual outcomes^21,78^.

Furthermore, while we focus here on behavior that is conserved across species (due to the limited number of studies directly comparing information seeking across species^8,88,89^ with data rich enough to test mathematically defined models of value), Info preference could vary across species based on their phylogeny and ability to gather and use information in nature. For example, while information seeking has largely been studied in terrestrial animals, work on the evolution of cognition suggests that opportunities to use predictive information may be limited in certain aquatic animals^64^; conversely, they may be expanded in certain avians, due their combination of visual acuity to detect distant rewards and flight to rapidly approach them. Indeed, recent reports indicate that goldfish may not prefer Info despite preferring rewards and learning cue-reward associations in a similar way to other species^90,91^ whereas pigeons have relatively strong Info preferences^88,89^, consistent with information value varying across species (Fig 1B). Our framework will enable future work to quantitatively determine the relative importance of the factors governing subjective value, how they vary across species, and which portions of this value could originate from specific advantages in naturalistic environments.

In addition, psychiatric disorders associated with aberrant information seeking strategies are prevalent in the modern “information age”^92,93^. This raises the possibility that maladaptive TA or CA-related processes may contribute to these conditions. Attention deficit disorders disrupt the ability to stay on task vs. leave it for other pursuits^94^ and are associated with altered stay-leave decisions during foraging^95^, while compulsive disorders can disrupt the proper assignment of credit for actual and possible outcomes^96,97^. Our framework may be able to provide insight into the structure of variation in information seeking strategies across individuals, which may be adaptive in natural environments with diverse information sources and computational problems to solve, but maladaptive and require adjustment in certain modern environments.

We note that theories on the origin of subjective value aim to explain behavior at a different level than conventional process models, so they also have different limitations^38^. Here we detail salient limitations and the necessary steps we took to handle them.

1. In this initial work we necessarily implement theories with naturalistic models that are simple and compact. This is to capture the theorized mechanisms without making strong assumptions about the detailed structure of environments, and to make it possible to derive optimal Bayesian agents for all theories to evaluate them comparably. As methods are developed to measure information landscapes in natural environments, it may become possible to formulate species-specific theories^64,98^, and test them with newly developed methods to compare subjective value across species^8,89^.
2. We apply our framework to information seeking about the size of uncertain rewards, because this is the information preference existing theories have been proposed to explain^6,37^ and where cross-species SVOIexp measurements are available to evaluate them^8^. Other information preferences may have distinct origins. Indeed, there is evidence that information seeking about rewards is distinct from information preferences about aversive events^99^ and information aversions^27,32,100^, which are encoded by partially distinct cortical circuits^101^, and may vary much more across individuals^101,102^.
3. Even if a theory correctly pinpoints the computational problem that subjective values originated to solve, this does not guarantee the values of real organisms solve them perfectly. For example, if information value originated through evolution by natural selection, then just like any other stochastic optimization process, it may not find the global optimum due to its inherent constraints (e.g. neural circuit structure, selection dynamics, and genetic and developmental constraints, etc.^103^). Therefore, we focus on theoretical predictions about core motivational factors that have high potential to be tuned by evolution or developmental learning: factors that strongly influence motivated behavior consistently across species and are strongly encoded by neural circuits. This includes expected reward, reward uncertainty, and cue and reward timing^1^.
4. Crucially, we test theories with subjective values measured in non-instrumental tasks. This is because preferences in instrumental tasks (e.g. where Info gives more reward than Noinfo) could occur simply through standard reinforcement learning and decision processes, without requiring any additional subjective valuation of information^2,6^.

Finally, we emphasize that our framework accommodates multiple potential origins of subjective value. Organisms can adapt to their environments in multiple ways, including evolution through natural selection over generations and developmental learning during an individual’s lifetime. In the case of information seeking, both are likely to contribute to its subjective value. Information value is likely to have a strong evolutionary component. It occurs across species that have very different developmental durations and experiences, and stubbornly persists despite long-term experience in non-instrumental tasks where information comes at a high cost^8,11^, suggesting that it is a conserved behavior that cannot be explained purely by simple reinforcement learning from experience. However, developmental processes are also likely to contribute to information seeking and motivation to resolve uncertainty^48,104^, especially in humans, other primates, and other organisms with long lifespans^105^. These origins may also be entwined. This is known to occur in neural systems as diverse as primary sensation, birdsong, and human language, which have all evolved to refine their circuitry during development through specialized forms of experience-dependent learning^106^. Thus, it is possible that neural systems for information value are implemented by evolved circuitry that is further refined by specialized forms of information-oriented learning during development. Our framework will enable future work to uncover how and why these value computations occur across species and individuals, and illustrates how the apparent paradox of “subjective value without instrumental value” can emerge as a natural consequence of organisms adapting to solve the computational problems posed by their environments.

## AUTHOR CONTRIBUTIONS

ESBM and JM conceptualized the project and theoretical framework. JM developed, implemented, and validated initial versions of the models. ESBM refined the framework, theories, and models, implemented and validated their final versions, and analyzed the data, in consultation with IEM. ESBM and IEM wrote the manuscript in consultation with JM.

## ACKNOWLEDGEMENTS

This work was supported by the National Institute of Mental Health under award numbers R01MH128344, R01MH110594, and R01MH116937; and the Conte Center on the Neurocircuitry of OCD MH10643 (to I.E.M.). We thank Dr. Jennifer J. Bussell and Dr. Victor Ajuwon for generously providing primary data from information choice tasks in mice and goldfish, respectively.

## METHODS

### General procedures

We analyzed experimental data from humans and monkeys performing analogous multi-attribute information choice tasks which have been described previously^8^ (Fig S6). All human procedures were approved by the Washington University Institutional Review Board. In total, 824 human participants completed tasks using Amazon’s Mechanical Turk service (mean self-reported age = 36.06 years, SD = 11.15 years; self-reported 410 female, 406 male, 4, other, 4 no response). All provided informed consent. Participants were monetarily compensated based on their performance, as detailed below. Participants were required to be healthy adults between the ages of 18 and 55 years with no history of previous neurological or psychiatric illnesses, located in the United States and able to read English and have normal or corrected-to-normal visual acuity. All animal procedures conformed to the Guide for the Care and Use of Laboratory Animals and were approved by the Washington University Institutional Animal Care and Use Committee. Four adult male rhesus monkeys (*Macaca mulatta*) participated in the experiments (animals R, Z, B and P; all 7–9 years old during the experiments).

### Information choice tasks for humans

In brief, humans performed tasks on the mTurk platform using mouse clicks to get monetary reward. The task had two versions, version 1 with Info vs. Noinfo (Fig 3) and version 2 with Early Info vs Late Info (Fig 4). Based on previous work, we analyzed the subset of participants who correctly answered a post-task questionnaire confirming they understood the stimulus feature that indicated an offer’s informativeness and that the information was non-instrumental, and whose choices could be validly fit by the GLM used for our analysis such that the fitting procedure converged and all weights were identifiable (e.g. their choices could not be entirely explained by a single bias like always choosing the offer on the left side of the screen). This resulted in a dataset of n=374/580 participants from version 1 and n=147/244 participants from version 2. Similar patterns of behavior were found if analyzing all participants, and regardless of self-reported age and gender^8^.

In version 1 (Fig S6), participants performed 150 trials, earning an average of $9.81 (s.d. = $0.41) from the entire task. Each trial started with a fixation point at the center of the screen which participants were required to click. Two offers appeared on the left and right sides of the screen. Each offer’s color indicated whether it was an Info or Noinfo offer, and a set of bars indicated its reward distribution. Each offer had four bars, each showing a stack of coins, representing four equally probable reward outcomes (1 coin = 1¢). The participant clicked their preferred offer, after which the non-chosen offer disappeared, and the trial proceeded with their chosen offer. If they chose an Info offer, they were shown an informative cue for ∼3.7 s indicating which of the four possible outcomes they would receive from the current trial. If they chose a Noinfo offer, they were not shown an informative cue for ∼3.7 s. Participants were instructed that information was non-instrumental and that the trial’s reward outcome would be the same and would be added to their winnings regardless of whether they chose Info or Noinfo.

Each offer’s informativeness and reward distribution was set randomly, as follows. The expected reward could be 5, 6, or 7 coins. The uncertainty of offers could be Safe, 25/50/25 or 50/50, where big vs. small reward outcomes were 4 coins above vs. below the offer’s expected reward. On 75% of trials the two offers were generated independently; on the remainder they were constrained to have different informativeness and identical reward distributions. To control for color preferences, the Info and Noinfo colors were counterbalanced across participants and reversed approximately halfway through the task; participants reported the identity of the Info and Noinfo colors for 10¢ bonuses on four ‘question trials’ distributed during the task. To control for side preferences, the left/right locations of the offers were randomized each trial.

In version 2 (Fig S6), participants performed 100 trials, earning an average of $6.45 (s.d. = $0.45) from the entire task. The format was similar but with the following key changes. All offers were 50/50. The cue period was longer in duration (∼14 s). Instead of Info and Noinfo, offers were Early Info and Late Info. Early Info offers showed an informative cue for the first ∼12 seconds of the cue period, then a second informative cue for the last ∼2 seconds. Late Info offers showed a non-informative cue for the first ∼12 seconds of the cue period, followed by an informative cue for the last ∼2 seconds.

### Information choice tasks for monkeys

In brief, monkeys performed similar tasks to humans, but in a lab environment, using eye movements to fixate and make choices, and receiving juice rewards. As before, offers were either Info or Noinfo indicated by their colors, and provided rewards drawn from a distribution shown by four bars indicating four equally probable reward sizes. Info offers showed an informative cue indicating which of the four rewards would be delivered, while Noinfo offers showed a non-informative cue that did not. To ensure animals had ample opportunity to prepare and receive the juice on every trial regardless of whether they chose Info or Noinfo, the reward size was always fully revealed by a ‘reveal’ stimulus a fixed time before juice delivery, and the juice spout was placed directly in the mouth.

In version 1 (Fig S6, n=4), offers had fixed times between events occurring after the choice: *T_choice-cue_* = 0.5 s, *T_cue-reveal_* = 3.5 s, *T_reveal-outcome_* = 0.5 s, *T_iti_* = 1.2 s. Offers varied in E[r], SD[r], and the type of reward distribution, including Safe, 25/50/25, and 50/50 offer types. As described previously, the precise settings of the reward distributions (e.g. maximum reward size, variation of expected reward across the offers, and ranges of reward used to construct distributions) varied slightly across animals^8^.

In version 2 (Fig S6, n=3), offers had variable cue and reveal/outcome times, indicated by a new “clock” added at the bottom of the offer stimulus. If the offer was chosen the clock then played an animation tracking the passage of time over the 8 s from the time of choice until the end of the trial. *T_reveal-outcome_* was fixed at 1.1 s. For each offer, the pair of event times (*t_cue_* and *t_reveal_*) was drawn uniformly at random from the set of all possible pairs meeting these requirements: *T_choice-cue_* between 0.4 and 5.8 s, *T_choice-reveal_* between 1.2 and 6.7 s, *t_cue_* at least 0.8 s before *t_reveal_*, and *t_cue_* and *t_reveal_* both multiples of 0.1 s. As described previously, this version also used slightly different reward distributions, e.g. by scaling up the reward sizes to compensate the animals for the longer trial durations^8^.

### Time Allocation model of foraging

The TA agent aims to maximize reward in an environment representing the process of pursuing a potential future reward outcome. The key intuition is that information helps the agent optimize stay-leave decisions: decisions about whether it will get a higher reward rate by *staying* in the pursuit or *leaving* to start another pursuit^37^. We model this with a hybrid approach combining elements of optimal foraging theory and reinforcement learning theory.

As in standard reinforcement learning, we model the environment as a Markov decision process (MDP) in which the agent is repeatedly confronted with a state *s* of the environment, then chooses an action *a* from a set of available actions in that state, then probabilistically transitions to a new state *s’* and receives a reward *r*, drawn from a distribution p(*s’*, *r* | *s*, *a*). We denote the agent’s policy for choosing actions in each state as π = p(*a* | *s*), and denote the optimal policy as π*. Each pursuit starts in a ‘start’ state and ends in an ‘end’ state. The MDP is designed to mimic the concept of the time allocation theory (Fig 2A,B) while also closely matching the temporal structure of experimental tasks to allow comparison with behavior, as follows.

We first describe the non-instrumental version of the environment (Fig 2D, left). Unless otherwise noted, each state has only one available action, to continue to the next state in the sequence. The start state leads to a ‘choice’ state lasting for duration *T_choice-cue_* with two possible actions, ‘choose Info’ and ‘choose Noinfo’. Choosing Info leads to one of several informative cue states. Specifically, we denote the pursuit’s initial reward probability as drawn from a distribution p(*r*) with *n* possible reward outcomes *r_1_,…,r_n_* each occurring with probability *p_1_,…,p_n_*. Choosing Info leads with probability *p_i_* to state ‘info cue *i*’. Each info cue state lasts for *T_cue-reveal_* and then leads to a corresponding ‘reveal state *i*’; that reveal state then lasts for *T_reveal-outcome_*, delivers the corresponding reward outcome *r_i_*, and transitions to an ‘outcome’ state lasting for *T_outcome-travel_*. Finally, the outcome state transitions to a ‘travel’ state lasting for *T_travel_* and then to a final ‘end’ state, representing traveling to a new pursuit. Choosing Noinfo leads instead to a non-informative cue state, which also lasts for *T_cue-reveal_* and then transitions with probability *p_i_* to each of the corresponding ‘reveal state *i*’. Thus the difference between Info and Noinfo is that Info leads to an informative cue state that reveals the future reward size immediately, while Noinfo leads to a non-informative cue state that means the agent has to wait *T_cue-reveal_* (i.e. *Tadvance*) more seconds until the outcome is delivered.

To extend the model to the leavable information task in which information is instrumental, every state is augmented with an additional action ‘leave’, which immediately transitions to the ‘travel’ state. This transition occurs at the start of the state where the action is taken, without waiting for the normal time duration of the current state to elapse. For example, if the agent is shown an informative cue giving ‘bad news’ and chooses to leave, they immediately move to the travel state, without waiting for *T_cue-reveal_* to elapse. We note that the agent could in principle choose to leave at any time within the state, but leaving immediately is strictly better in all environments we examine here, because leaving later would simply waste time without any benefit.

As in optimal foraging theory, we optimize the agent’s performance in terms of the average reward rate, defined as the expected future reward per unit time; by contrast to standard reinforcement learning which optimizes time-discounted future reward based on a pre-specified discount factor^40^. To compute this tractably, for events occurring at a specified time *t* after the start of the pursuit, we define state-action values V_π_(*s*,*a,t*) as the average reward rate obtained starting from that state-action pair, then following policy π for a time span from *t* to a fixed maximum time point *t_max_* = 20 s. The pursuit itself is considered to last from time *t* = 0 until the travel state transitions into the end state at *t_end_*. All excess time after that point – that is, the time interval between *t_end_* and *t_max_* – is credited as producing rewards at a rate equal to the environment’s average reward rate *RRavg*(π). This captures the intuition from optimal foraging theory^63^ that leaving a pursuit early means the agent abandons the opportunity to gain the pursued reward, but in exchange, ends the pursuit sooner and can use the saved time to gain rewards elsewhere in the environment at the rate specified by *RRavg*(π). To compute values, we define R_π_(*s,a,t*) and T_π_(*s,a,t*) as the expected reward to be collected during the pursuit, and the expected time *t_complete_* when the pursuit will be completed. Thus, we define state-action values as:

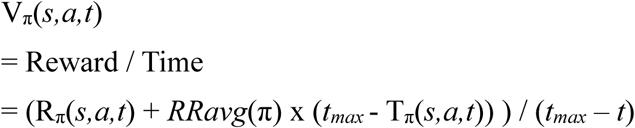

To jointly compute π*, V_π*_, and *RRavg*(π*), we use the following procedure. We first initialize V(*s,a,t*) = 0 and *RRavg* = 0 and initialize T_π_(*s,a,t*) based on the time duration parameters. We then repeat the following sequence of three steps for n=100 iterations: (1) Update π by setting it to choose the action in each state that has the highest value. (2) Given that π and *RRavg,* update Rπ, Tπ, and Vπ based on a standard dynamic programming sweep through the space of (*s,a,t*), starting with later times. (3) Given that Vπ, update *RRavg* by setting it to the value of starting pursuit, max*_a_*(Vπ(*start,a,0*)). We find this process consistently converges to self-consistent π*, V_π*_, and *RRavg*(π*) for the environments studied here in less than 10-20 iterations. This procedure accurately estimates the reward rate obtained by an agent carrying out the optimal policy, as indicated by the close correspondence between the reward rates obtained from simulation vs. computed by this procedure (Fig 2D vs 2E). To aid in comparing the model’s predictions against experimental data where trials with different reward distributions and/or event timings are interleaved in the same lab environment, we also implemented a procedure to allow separate runs of the model with different reward and/or timing parameters to be optimized given the same setting of *RRavg*. This procedure is the same, except *RRavg* is initialized and fixed to a specified setting (*RRavg* = 0.115 for task version 1, 0.05 for task version 2 that had longer trial lengths). The latter procedure was used to generate the results shown here except the simulation panels of Fig 2 mentioned above, in order to make the model as comparable as possible to experimental data where offers with different reward statistics were interleaved, but the former procedure produced qualitatively similar results.

All simulations compared with behavioral data are made with parameter settings to mimic the reward distributions and time courses used in the animal behavior experiments analyzed here^8^. To mimic experiments with varying reward distributions (Fig 4), we consider 81×5 = 405 reward distributions, using all combinations of 81 settings of E[r] and 5 settings of distribution shape. The settings of E[r] ranged from 0.5 to 1.5. The settings of distribution shape were: Safe, 25/50/25 with range = 0.7, 25/50/25 with range = 1, 50/50 with range = 0.7, and 50/50 with range = 1. To mimic experiments focused on reward timing instead of distribution shapes (Fig 5), we 3×5 = 15 distributions, using 3 settings of E[r] (0.5, 1.0, and 1.5). For experiments manipulating reward distributions (Fig 2,4,6), the timing parameters are as follows. Durations (in seconds): *T_choice_* = 0.5, *T_choice-cue_* = 0.5, *T_cue-reveal_* = 3.5, *T_reveal-outcome_* = 0.5, *T_outcome-iti_* = 0, *T_iti_* = 2. For experiments manipulating timing (Fig 5), they are *T_choice_* = 0.5, *T_reveal-outcome_* = 1, *T_iti_* = 2, and the remaining parameters set by the 78 combinations of *T_choice-cue_*, *T_cue-reveal_*, and *T_outcome-iti_* obeying the following constraints: all three are multiples of 0.5, all three are ≥ 0.5, all three sum to 8. For clarity, all diagrams of state value over time (e.g. Fig 2C) are made from simulations with parameter settings based on the experiments manipulating distributions but altered to mimic the time course of the classic information seeking task with no reveal stimulus (Fig 1A): *T_choice_* = 0, *T_choice-cue_* = 1, *T_reveal-outcome_* = 0.

To derive the minimal partial information the TA model requires to maximize reward rate, we note that the only actions the model makes that can influence its reward rate are its stay-leave decisions, which it makes at each moment by comparing the expected future reward rate from staying in the current pursuit versus the expected future reward rate from leaving the pursuit. Thus, partial information about the binary result of this comparison is sufficient for the model to achieve its maximal performance, producing the same reward rate as the standard full information (i.e. indicating the exact future reward magnitude and timing), as validated by our simulations (Fig 4). For comparison between TA and CA models using partial information (Fig 4), we use a reward distribution that is mixture of Gaussians that draws the outcome with equal probability from a ‘good’ N(+1, σ_noise_^2^) distribution or a ‘bad’ N(-1, σ_noise_^2^) distribution, with σ_noise_ = 1; discretizing the reward distribution into 22 equally spaced reward amounts from -5.25 to +5.25; and set *RRavg* = 0.15 to approximately maximize IVOImodel when full information is available. Partial information about stay vs. leave is a binary indicator of whether the value of staying is higher than the value of leaving (by leading to an ‘info cue stay’ or ‘info cue leave’ state), while partial information about local good vs. bad is a binary indicator of whether the reward was drawn from the good or bad distribution (by leading to an ‘info cue good’ or ‘info cue bad’ state), neither of which indicates the exact reward size.

### Credit Assignment model of reinforcement learning

In this model, agents use information about future rewards to optimize credit assignment in dynamic reinforcement learning environment. In a given environment, an agent repeatedly takes an action *a*, then observes a cue *c*, then obtains a reward outcome *r*, respectively occurring at times 0, *t_cue_*, and *t_outcome_*. The agent’s goal is to maximize the reward rate of its actions, defined as E[*r*] per action. The total reward from an action equal to the sum of contributions from the chosen action, *r_a_*, and from background events, *r_bg_*. Thus:

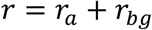

The agent has a binary choice between two actions, *a_1_* and *a_2_*. At any time one of these actions is the ‘good’ action a*, contributing a reward drawn from the distribution *p*(*r_a_* | *a* = *a\**) with mean E[*r_a_* | *a* = *a\**] = *r\**. The other is the ‘bad’ action *a*^-^, which contributes an opposite reward distribution *p*(*r_a_* | *a* ≠ *a\**) = *p*(-*r_a_* | *a* = *a\**).

The agent is not told which action is correct so it has to learn from experience. There are three obstacles the agent must overcome. First, the environment is dynamic: after each outcome, the correct and wrong actions randomly switch with probability *p_volatility_*. Second, the actions produce variable rewards: draws from *p*(*r_a_* | *a*) are equal to its mean E[*r_a_* | *a*] plus a draw ε from a noise distribution. In different environments shown here, the noise distribution is either a Gaussian (ε ∼ N(0, *σ_a_*^2^)), or is modeled after the discrete distributions from experimental data (Safe: 100% chance of ε = 0; 50/50 distribution with range *z* = 0.7 or 1.0: 50% chances of ε = -*z* or +*z;* 25/50/25 distribution with *z* = 0.7 or 1.0: 50% chance of ε = 0 and 25% chances ε = -*z* or +*z*). All discrete distributions are then convolved with a smoothing Gaussian N(0, 0.1^2^) to produce continuous reward distributions. Third, the background also produces variable rewards: *r_bg_* ∼ N(0, *σ_bg_*^2^).

The agent can use information to improve its credit assignment because, as in natural environments, the agent’s action and background contributions to the reward are modeled as unfolding over time. To represent this, we define V*(t)* as a continuous variable indicating the current state value, in terms of the total reward that has been scheduled so far to be delivered as part of the outcome, considering all events occurring from the start of time (time -∞) up to the present moment (time *t*). The cue c is either an informative cue that indicates the reward scheduled so far (Info condition, *c* = V*(t_cue_)*) or a non-informative cue (Noinfo condition, *c* = 0). The action’s contribution to the outcome occurs in a discrete manner, incrementing V by *r_a_* at time 0. The background’s contribution to reward is modeled as arising from a noise process *x_bg_*(*t*) that unfolds over time in time steps *Δt*, at each time step drawing from a Gaussian with mean = 0 and variance:

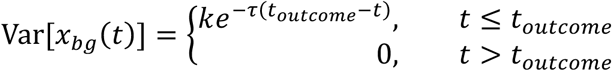

…where the timescale of background contributions to the reward is set by τ. We note that, while we chose this exponential function for simplicity, the same pattern of results would result as long as the time course is a monotonic function that changes gradually with time (e.g. events at some arbitrary specific time point do not have a special, privileged causal influence on the reward).

The key requirement is that the background needs to make a consistent contribution to the reward that unfolds gradually over time, so that graded changes in cue timing produce graded changes in learning performance. In principle the same results would hold even if this was a temporally flat function of *t* or a decreasing function of *t*, at least over the timescales considered in our experiments. The drawback of such flat or decreasing functions is that they cannot hold as a general rule over long timescales (because they would imply that events that are increasingly temporally distant from a reward still make either the same contribution or make increasingly large contributions to that reward; and hence the summed contribution of all past time points to the reward would have infinite variance). The constant *k* is chosen so the variance of the summed background contributions over all time before the reward delivery achieve the desired setting of *σ_bg_* for the environment, that is:

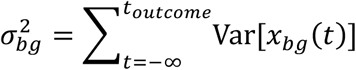

Crucially, an agent with access to Info can improve learning by using the cue to perform better credit assignment – more accurately assigning credit to the action for the portion of the reward it was responsible for producing. To see this, note that we can define the background reward as the sum of its pre-cue portion (from *t* = -∞ to *t_cue_*) and post-cue portion (from *t_cue_* to *t_outcome)_*, and hence its variance as the sum of their variances:

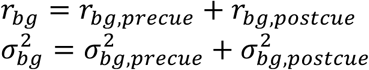

An agent in the Noinfo condition has to learn from *r* as a teaching signal, which includes the contribution of the action *r_a_* plus the full contribution of the background noise, whereas as agent in the Info condition can learn from *c* as a teaching signal, which includes only the pre-cue portion of the background noise:

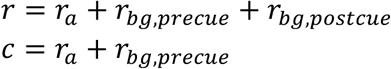

…effectively reducing the variance of the teaching signal by 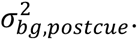 We used parameters *r\** = +1, *p_volatility_* = 0.1, τ = 0.2; other settings produced a similar pattern of results.

To implement an optimal Bayesian agent for this model, we use the standard approach of deriving a recursive Bayesian estimator for the key variable for action selection in this environment: the good action a*. We define prior_i_(a*=a) as the agent’s prior about a* before making its i-th choice in the environment. After the agent makes its choice and observes the resulting feedback (c for the Info agent, r for the Noinfo agent), it uses the likelihood of that feedback to convert its prior to a posterior using Bayes rule:

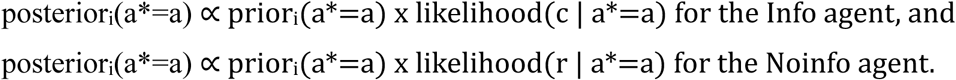

Finally, the agent updates this posterior belief about a* for the i-th choice to become a prior belief about a* for its upcoming i+1-th choice. This accounts for the possibility that a* changed in between the i-th and i+1-th choices, based on the volatility of the environment:

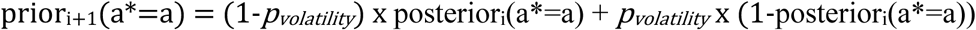

Importantly, this model environment is designed to directly measure information’s benefit for learning in a manner that is independent of the agent’s policy and action selection. That is, as long as the agent learns using the Bayesian estimator described above, the agent always receives the same quality of feedback for updating its beliefs regardless of its policy and regardless of whether any given choice it makes is good or bad. This is because both good and bad actions provide equivalent amounts of evidence in terms of the magnitude of the likelihood (as long as their reward distributions are symmetric to each other around 0).

To compute IVOImodel for a given set of parameters, we generate *n_env_* = 80 simulated environments using those parameters each lasting for *n_trials_* = 1000 trials, record the performance of an optimal agent in these environments given either informative cues or non-informative cues, then compute IVOImodel as the difference between their reward rates. For Fig 2 we use σ_a_ = 0.5 and σ_bg_ = 1; for Fig 3 we use the same parameters for the CA model as the TA model for properties of reward distributions, information timing, and reward timing, and σ_bg_ = 0.5; for Fig 4 we use σ_a_ = 1 and σ_bg_ = 1.5; Figs 2–3 use σ_a_ set to match the distributions used in the TA model and σ_bg_ = 0.5. Other parameters produced qualitatively similar results. Note that E[r] has no effect on this model because it simply adds an additive offset to all reward deliveries and hence does not alter learning; and *t_cue_* and *t_reveal_* only effect this model through their joint effect of governing *T_advance_*, not separately.

To derive the minimal partial information the CA model requires to maximize reward rate, we note that after the model makes choice *i*, the only way the model uses feedback from its observations (including both its observation of the cue, and later, its observation of the actual outcome) is to update its posterior belief about a*. That is, to update its belief about the underlying action values: which actions were good vs. bad at the time of its choice. Thus, partial information that produces the same distribution of belief updates (conditioned on its choice) is sufficient for the model to achieve its maximal performance, producing the same reward rate as the standard full information, as validated by our simulations (Fig 4). For example, if the model agent is not provided with the actual cue and reward observations drawn from *p*(*c*, *r* | *a*, *a\**), and instead is provided with a fresh random draw of hypothetical *c* and *r* from that distribution, then this would result in the same statistical distribution of updated posterior belief distributions conditioned on its choices, and hence statistically indistinguishable learning performance. For comparison between TA and CA models using partial information (Fig 4), we implemented agents that use optimal Bayesian inference to update their beliefs about action values based on observing both a partially informative cue *c_partial_* and the final reward outcome *r*. Partial information about stay vs. leave is a binary indicator of whether that current state value is higher or lower than a threshold θ representing the average reward rate *RRavg*. To obtain results with θ = *RRavg*, we initialize θ = 0.3, then iteratively repeat these two steps until θ converges to a fixed value: (1) run the simulation, (2) set θ equal to the empirical RRavg from that simulation. In practice we find this converges quickly (< 4 iterations). Partial information about local good vs. bad is a cue *c_partial_* ∼ N(E[r | V(*a*), σ_partial_^2^), where V(*a*) is the true expected value of the chosen action (*r\** for the good action, *-r\** for the bad action) and the setting of σ_partial_ is derived so an optimal Bayesian agent given both *c_partial_* and *r* gets a matched quality of evidence about V(*a*) as an agent given full information in the form of *c*, resulting in identical reward rates in a dynamic reinforcement learning environment.

### Mixed value computations

We model the information seeking performance of agents in a set of environments where the instrumental value of information arises from both time allocation and temporal credit assignment problems, and where agents can pay for different types of full or partial information to help solve these problems. Unlike the other agents we examine, these agents are not optimized based on any one specific model for any one specific environment (i.e. setting SVOI equal to IVOImodel computed by a single model for a single environment). Instead, these agents set their SVOI of partial information based on a mixture of the patterns of IVOImodel from the TA and CA models (Fig 5A). This represents a situation where a species adapts to environments where they must face both problems, and hence has subjective value computations that include elements of both TA and CA theories.

Here we first define the set of environments the agents face, and then define the agent’s subjective values of information. We consider a set of *n* environments *e*_1_, *e*_2_, …, *e_n_*. Each environment *e* has the same, fixed, total IVOI available as all other environments. However, each environment has a unique mixture fraction *m*(*e*) of its IVOI arise from information’s benefit for time allocation (TA), while the remaining (1-*m*(*e*)) is obtained from information’s benefit for credit assignment (CA). We set *n* = 11 with their corresponding *m*(e) = 0, 0.1, …, 0.9, 1.0. We model this in the simplest possible way: each environment presents the agent with a sequence of situations *s*, consisting of TA situations which occur with probability p(*s* = TA | *e*) = *m*(*e*) and CA situations which occur probability p(*s* = CA | *e*) = 1-*m*(*e*). Thus,

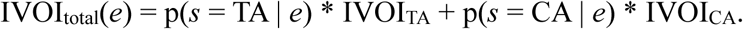

We then define the fraction of this total IVOI that each type of information provides for TA and CA situations, based on our modeling results using partial information. Specifically, we define IVOI(*s*,*i*) for each situation type s and information type *i*, where the three types of information are *i* = 1, 2, 3 for full information, partial information about local vs. global reward rates, and partial information about local good vs. bad states. To do this, we define the maximal IVOI = 1 as occurring when the agent receives full information (*i* = 1), or receives the appropriate type of partial information for its current situation (i.e. *i* = 2 for *s* = TA, or *i* = 3 for *s* = CA). Thus, IVOI(TA,1) = IVOI(CA,1) = IVOI(TA,2) = IVOI(CA,3) = 1. We define the partial IVOI as occurring when the agent receives the inappropriate type of partial information for its current situation (i.e. *i* = 2 for *s* = CA, or *i* = 3 for *s* = TA). Thus, we set IVOI(TA,3) = IVOI(CA,2) = 0.54. This is based on our modeling results finding that the inappropriate type of partial information provides ∼0.54 of the IVOI of full information in both TA and CA environments with the tested parameters (Fig 4C). We define the minimal IVOI = 0 as occurring when the agent receives no information.

Finally, to allow the agent to trade off information for primary reward, we allow the agent to pay for information. Specifically, in each situation, the agent is offered an opportunity to pay a random cost *c* for a random type of information. The cost *c* is drawn from a discrete uniform distribution from 0 to 2 in steps of 0.01. The type of information is drawn from a discrete uniform distribution over the three types of information, each occurring with equal probability (full, partial about local vs. global reward rates, partial about local good vs. bad states). If the agent chooses to pay the cost, the agent receives the information but loses *c* reward in exchange. If the agent chooses not to pay the cost, the agent receives no information.

We endow each agent *a* with a subjective value function SVOI*_a_*(*i*) that assigns subjective value to each of the three types of information. The agent uses this subjective value to make its decisions to pay for information. That is, the agent will pay a cost *c* for information *i* if and only if the agent’s SVOI*_a_*(*i*) > *c*. Thus, the information value obtained by the agent *a* in a given situation *s* from an offer of information type *i* for a cost *c* is:

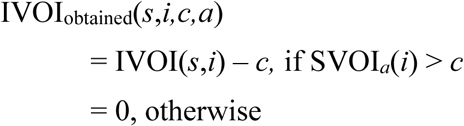

We can then simply calculate the IVOI the agent obtains in each environment, by summing its IVOI obtained from each possible combination of three variables that can occur in that environment (situation, offered information type, offered information cost), weighting each of them by its probability:

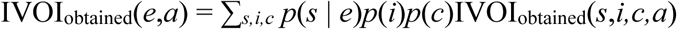

And use this to calculate the overall IVOI the agent obtains from the full set of environments, by weighting each environment by its probability (which uniform over the environments in the set; that is, p(*e* = *e_j_*) = 1/*n* for all *j*):

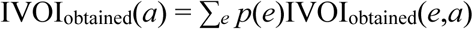

We tested endowing agents with several types of such value functions. First, value functions where SVOI(*i*) is a linear mixture of the true instrumental value of each type of information for TA and CA purposes, with a mixing factor *α*:

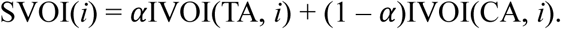

We tested value functions with 11 different settings of *α* = 0, 0.1, …, 1.0. Thus, an agent with *α* = 1 always values all types of information based only on their bene[it in TA situations, an agent with *α* = 0 always values all types of information based only on their bene[it to CA situations, and an agent with *α* = 0.5 values information based on 50/50 mixture of its bene[its for TA and CA situations. As controls, we also tested value functions that do not consider the differences between the different types of partial information. Speci[ically, value functions that set the SVOI(*i*) for all types of partial information (*i* = 2 or 3) equal to the maximum of all types of partial information, minimum of all types of partial information, or to zero. We also tested a value function that sets SVOI(*i*) = 0 for all types of information.

To infer an agent’s SVOI of partial information from its choice data, we simulated agents with SVOI of partial information (relative to full information) equal to 0, 0.025, …, 1.0. Each agent made n=10 choices to either pay or not pay for each type of information in each of the above situations in each environment, using a softmax choice rule with inverse temperature parameter equal to 1. For each environment, we inferred SVOI by minimizing the mean squared error between the predicted vs. actual choices (where the predicted choice is 1 (accept) if SVOI > *c* and 0 (reject) otherwise).

### Data analysis

All statistical tests are two-tailed unless otherwise noted, and analysis of behavior is done on successfully performed trials. To quantify the effects of motivational factors on subjective value (SVOIexp), we analyzed human and monkey choice data using the same analysis framework as in previous work^8^. In brief, choices from each individual were fit with a standard logistic GLM for binomial data:

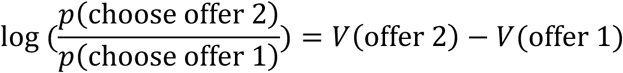

…where the value of each offer, V(offer), is a linear weighted combination of the offer *i*’s attributes *x_i,_*_1_, …, *x_i,n_*:

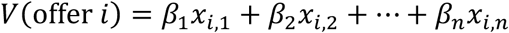

All non-indicator attributes were standardized before analysis to allow comparison of weights between variables with different units (e.g. ¢ vs ml juice). The resulting fitted β weights estimate effects as subjective values in units of log odds of altering choice. We then re-scaled them to be in units of reward (by simply dividing them by the fitted effect of 1¢ of E[r] for humans, or 1 ml juice of E[r] for monkeys). Thus, the estimated effect of a given factor on SVOIexp is the weight of the regressor “Info x (factor)”.

We used an analogous analysis to quantify the effects of motivational factors on instrumental value in the models (IVOImodel). For each TA model analysis, for each tested model environment (i.e. set of parameters specifying motivational factors), we defined the value of Info and Noinfo offers as their values according to the optimal policy, i.e. V_π*_(choice, choose Info*, t_choice_*) and V_π*_(choice, choose Noinfo*, t_choice_*). For each CA model analysis, for each of the same tested environments, we defined their values as the reward rate of the optimal policy when the cue indicating the reward outcome was provided at the time corresponding to the Info condition (i.e. *t_cue_*) or Noinfo condition (i.e. *t_reveal_*). We then calculated IVOImodel as the difference in value between Info and Noinfo. Finally, for each model and in each analysis, we fit all of its IVOImodels from all tested environments using a standard linear GLM for normal data:

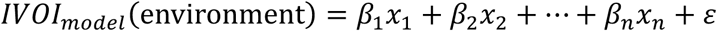

…where *x* are the corresponding attributes of that model environment and ε is a normally distributed error term. Thus, the estimated effect of a given factor on IVOImodel is simply the weight of that factor. For human data, we examined the mean fitted weights over all analyzed participants; for monkey and model data, we examined the fitted weights separately for each tested monkey or model. Finally, the regressors used for each GLM are shown in Table S2.

## SUPPLEMENTAL MATERIAL

**Figure S1.**
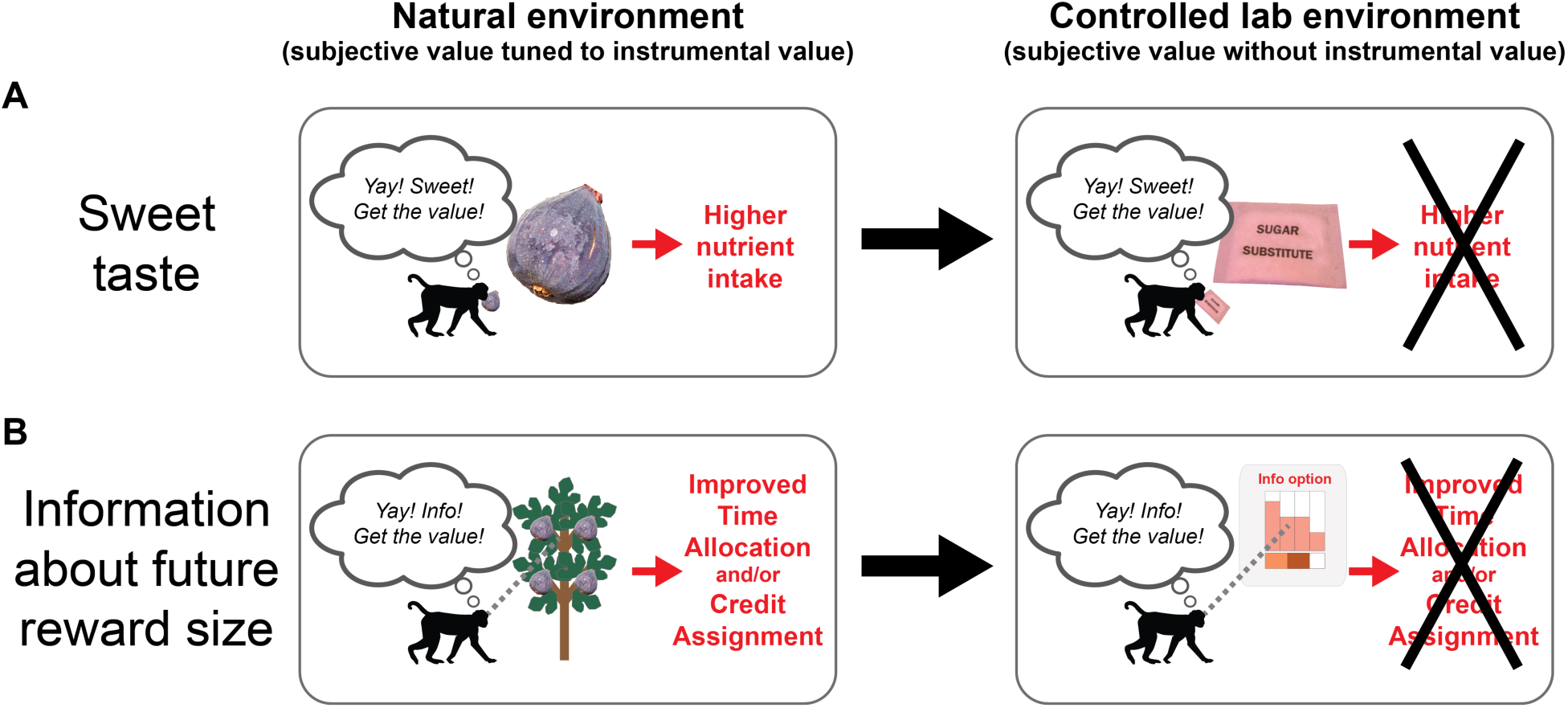
Theoretical framework for understanding the origin of subjective value without instrumental value. **(A)** It has been proposed that the subjective value of tastes evolved due to their instrumental value for motivating the behavior of organisms in natural environments. In this view, animals place positive value on sweet taste because of its instrumental value as a natural cue to sugar, and negative value on bitter taste because of its instrumental value as a natural cue to toxins^108^ (left). This explains why humans and many animals evolved to persistently pay for sweet taste, even if that sweet taste is gained from artificial sweeteners that are available in modern and/or laboratory environments, that have no nutrient content (right). **(B)** We and others have proposed^6,37^ that the same is the case for the subjective value of information about the size of future rewards. Similar information may have instrumental value for organisms to solve core computational problems posed by their natural environments, such as the Time Allocation problem and Credit Assignment problem in the theories discussed here (left). As a result, these organisms may have evolved to place subjective value on this information, even in modern environments or laboratory environments where it has no instrumental value because it cannot be used to control the outcome (right). While certain species may have evolved to place lower value on this information, potentially including goldfish^91^ (Fig 1B), perhaps due to them either having more limited abilities to gather and use this information in their natural environments, or due to their survival and reproduction not being as strongly affected by those computational problems.

**Figure S2.**
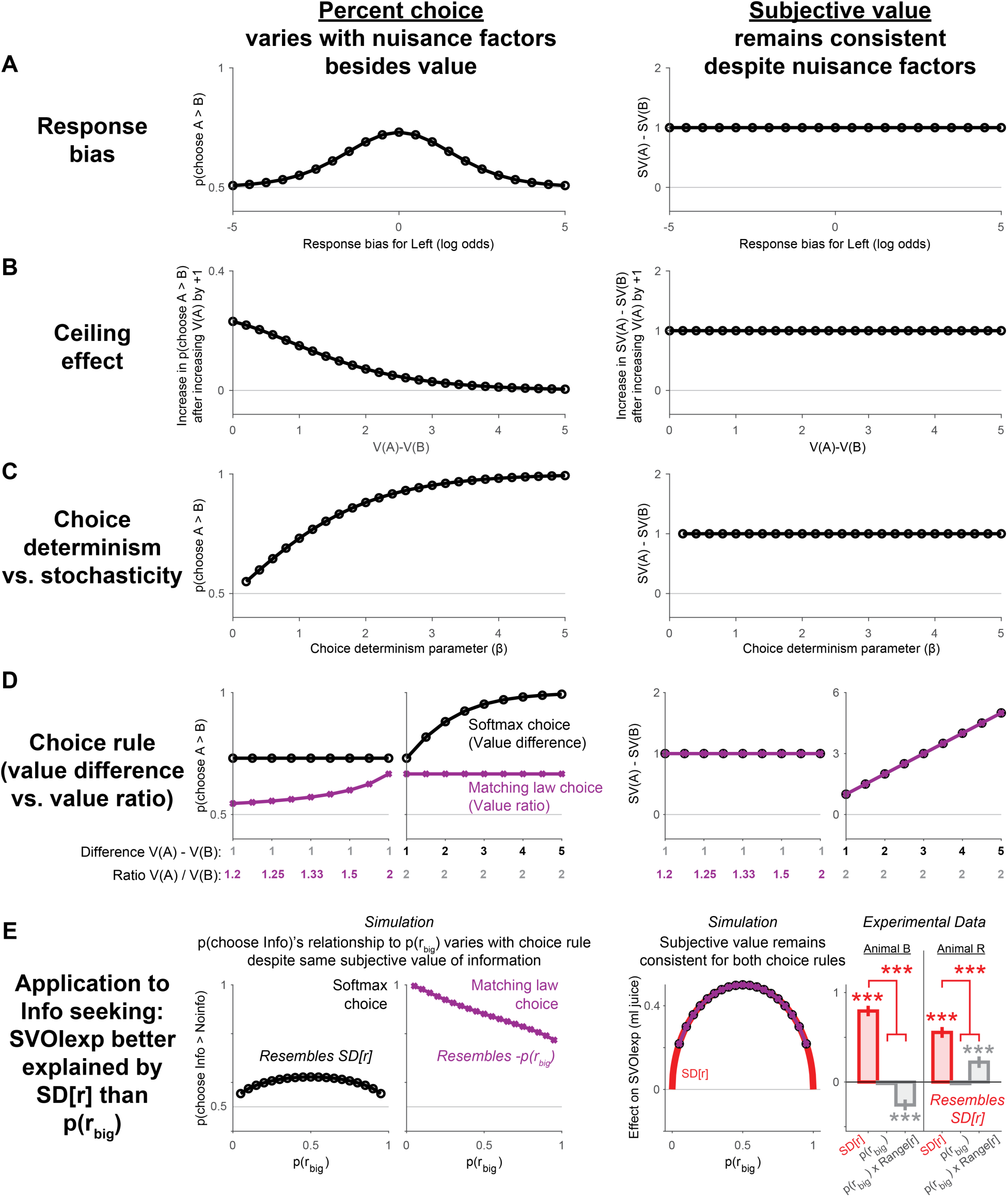
Understanding the origin of subjective value requires measurement of subjective value, not only percent choice. Here we show several examples demonstrating that percent choice may not faithfully track the subjective value that governs preference, due to confounding factors that may vary across conditions, individuals, and species. We use models following the common framework of value-based decisions in which an agent chooses between offer A and offer B based on their subjective values V(A) and V(B). For each scenario, the *Left* side shows how confounding variables cause percent choice to vary across conditions, while the *Right* side shows how subjective value does not vary in those manners. The agent translates values into choices using one of two popular choice functions used to model behavior: *softmax* (black, based on the difference of values: p(choose A)/p(choose B) = exp(β(V(A)-V(B))); β=1 unless otherwise noted) and the *matching law* (purple, based on the ratio of values: p(choose A)/p(choose B) = V(A)/V(B))^40,109^. **(A) Response bias.** *Left:* Percent choice of the preferred option A (with value V(A) = 2) over the non-preferred option B (with value V(B) = 1) varies greatly as a function of response bias favoring whichever option happens to be randomly presented on the left side of the screen on each trial (x axis, log odds). For example, if response bias is very strong then the bias entirely determines choice and so the subjective value of A vs B has no effect on percent choice. *Right:* subjective value difference remains constant regardless of response bias (V(A)-V(B) = 1). **(B) Ceiling effect.** *Left:* If the value of A is increased by a fixed amount (V(A’) = V(A) + 1), the increase in percent choice varies greatly as a function of the pre-existing value difference (V(A)-V(B), x-axis). For example, if V(A)-V(B) = 5, then the agent already chooses A 100% of the time, so increasing its value further has no effect on choice. *Right:* increase in subjective value remains constant regardless. **(C) Choice determinism vs. stochasticity.** *Left:* popular choice rules have an adjustable level of stochasticity vs. determinism, governed by a determinism parameter β (which may vary widely across conditions, individuals, and species). This controls how values are translated into precent choice. For the softmax rule used here, choice is random with β = 0 (50% choice of A), and becomes nearly deterministic with β = 5 (100% choice of A). *Right:* subjective value difference remains constant regardless. **(D) Choice rule.** *Left:* the popular *softmax* choice rule sets percent choice based on the value *difference* (V(A)-V(B)). However, another popular choice rule used to fit descriptive models of behavior is the *matching law*, which sets percent choice based on the value *ratio* (V(A)/V(B)). Thus, individuals using a softmax rule (black) will show percent choice that is constant for a constant value difference (left) but varies despite a constant value ratio (right). Individuals using a matching law rule (purple) will show percent choice with the reverse pattern, varying for a constant value difference but not a constant value ratio. *Right:* measured subjective value difference between the options, in terms of willingness to pay, accurately recovers the subjective value difference regardless of the individual’s choice rule (black = purple). **(E) Measuring subjective value with our data shows that info preference can be better explained by SD[reward] than p(big reward), unlike what has been assumed based on measuring only percent choice**. Many previous studies reported the percent choice of Info to be negatively related to the probability of reward p(reward), or to probability scaled with reward magnitude (e.g. p(big reward) x Range[reward]). For example, pigeons and rats have been reported to have a higher percent choice of Info for p(reward) = 25% than p(reward) = 75%^27^. This was interpreted as evidence against the hypothesis that information preferences are governed by symmetric uncertainty measures (like SD and Shannon entropy) which have very different functional forms as inverted-U functions of p(reward)^110, 111^ (Right, red curve). *Left:* However, this interpretation may be incorrect due to the confounds discussed above. Consider the simplest possible scenario in which animals assign Value(offer) = Value(reward) + Value(information) = E[r] + SD[r]. The animals percent choice would only have an inverted-U shape if they used a softmax choice rule (black). Their percent choice would instead have a strongly negative shape, similar to previously reported data, if animals instead used the matching law choice rule that is commonly used to model the behavior of those species in similar experiments (purple). This is because the matching law says a fixed increase in Value(information) causes a greater increase in percent choice when both options have low Value(reward). For example, if p(reward) = 25% then percent choice of Info is 94%, but if p(reward) = 75% then percent choice of Info is 83% (former case: V(Info) = 0.68, V(Noinfo) = 0.25, ratio = 2.73; latter case: V(Info) = 1.18, V(Noinfo) =0.75, ratio = 1.58). We also note that manipulating p(reward) over most of its range is only a surprisingly modest manipulation of SD[r] (varing p(reward) from 0.25-0.75 varies SD[r] only from 0.43-0.50), so manipulating the sizes of the reward outcomes would provide a stronger test. *Right:* Measuring the subjective value of information (SVOIexp) can recover subjective value regardless of the choice rule (black line = purple line) and can thus test whether it is governed by a negative effect of p(reward) or a positive effect of SD[r] (red line). We therefore tested this in our experimental data. For two monkeys, the set of reward distributions in some experiments included a subset of offers giving p(big reward) = 1 – p(small reward) = 25%, 50% or 75%, with varying big reward sizes *r_big_* and small reward sizes *r_small_* (animal B, n=23271/42076 offers from this subset; animal R, n=11174/20590 offers from this subset)^8^. This made it possible to measure how SVOIexp was affected by the key motivational factors discussed here: SD[r], p(big reward), and p(big reward) x Range[reward]. To do this, we extended our GLM from Fig 4A to include separate weights for each of the these three key factors to modulate SVOI for that subset of offers. Shown are fitted beta weights, in units of ml juice, of non-normalized regressors for the effects on SVOIexp of SD[r] (red bar), as well as p(big reward) and p(big reward) x Range[*r*] (gray bars). Both animals were best fit by very large positive effects of SD[r], near-zero effects of p(big reward), and smaller and idiosyncratically-signed effects of p(big reward) x Range[*r*] (negative for one animal, positive for the other). SD[r] had a significantly greater effect than both p(big reward)-based factors in both monkeys (p < 0.006 for SD[r] vs each other bar; 2-sided t-test of GLM weights).

**Figure S3.**
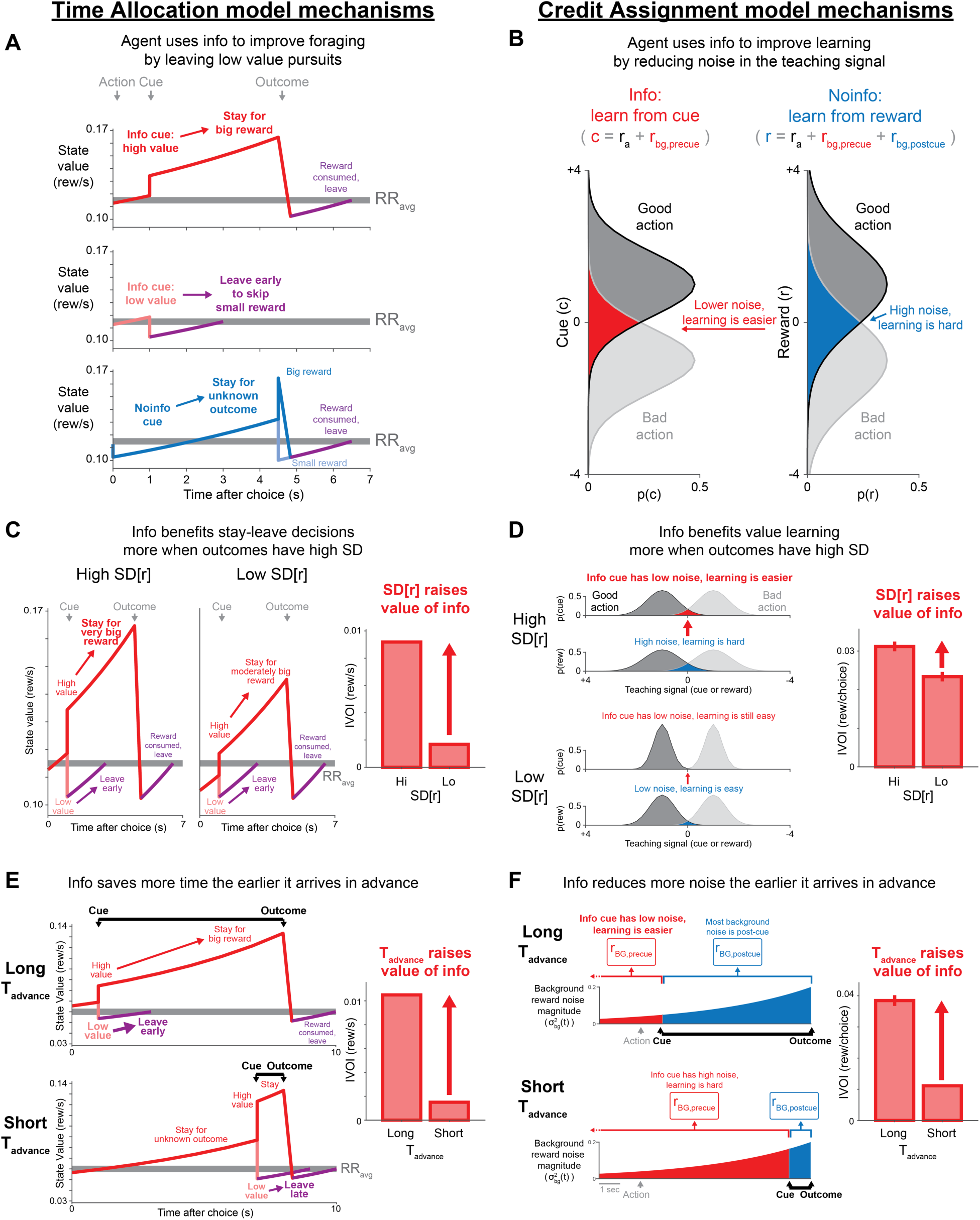
Mechanisms of information value computations in TA and CA models. Left: TA model. Right: CA model. **(A)** TA model mechanism of instrumental value: leaving early from low-value pursuits. State value is shown over time for Info→big or small reward (top, red; middle, light red), and Noinfo→big or small reward (bottom, blue and light blue); also after leave actions (purple) and *RRavg* (gray). **(B)** CA model mechanism of instrumental value: agents learn better from the cue since it is only affected by a fraction of the background noise (*r_bg,precue_*, red; not *r_bg,postcue_*, blue). Credit assignment improves because good and bad actions are more distinguishable (red vs. blue shaded areas). **(C)** TA model mechanism of information value scaling with SD[r]. Agents gain more from Info for offers with high than low SD[r] (left vs. middle), because on average they have larger reward rate difference from staying vs. leaving (cued state value vs. *RRavg*), producing higher IVOI (right). **(D)** CA model mechanism of information value scaling with SD[r]. Agents gain more from Info for offers with high than low SD[r] (top vs. bottom), because higher noise makes learning hard and Info reduces that noise, producing higher IVOI (right). **(E)** TA mechanism of information value scaling with Tadvance. Agents gain more from Info with long Tadvance because they save more time leaving early (top) than late (bottom), producing higher IVOI (right). **(F)** CA mechanism of information value scaling with Tadvance. Agents gain more from Info with long Tadvance because it reduces pre-cue background noise (red, top vs. bottom) which makes cues better teaching signals, producing higher IVOI (right).

**Figure S4.**
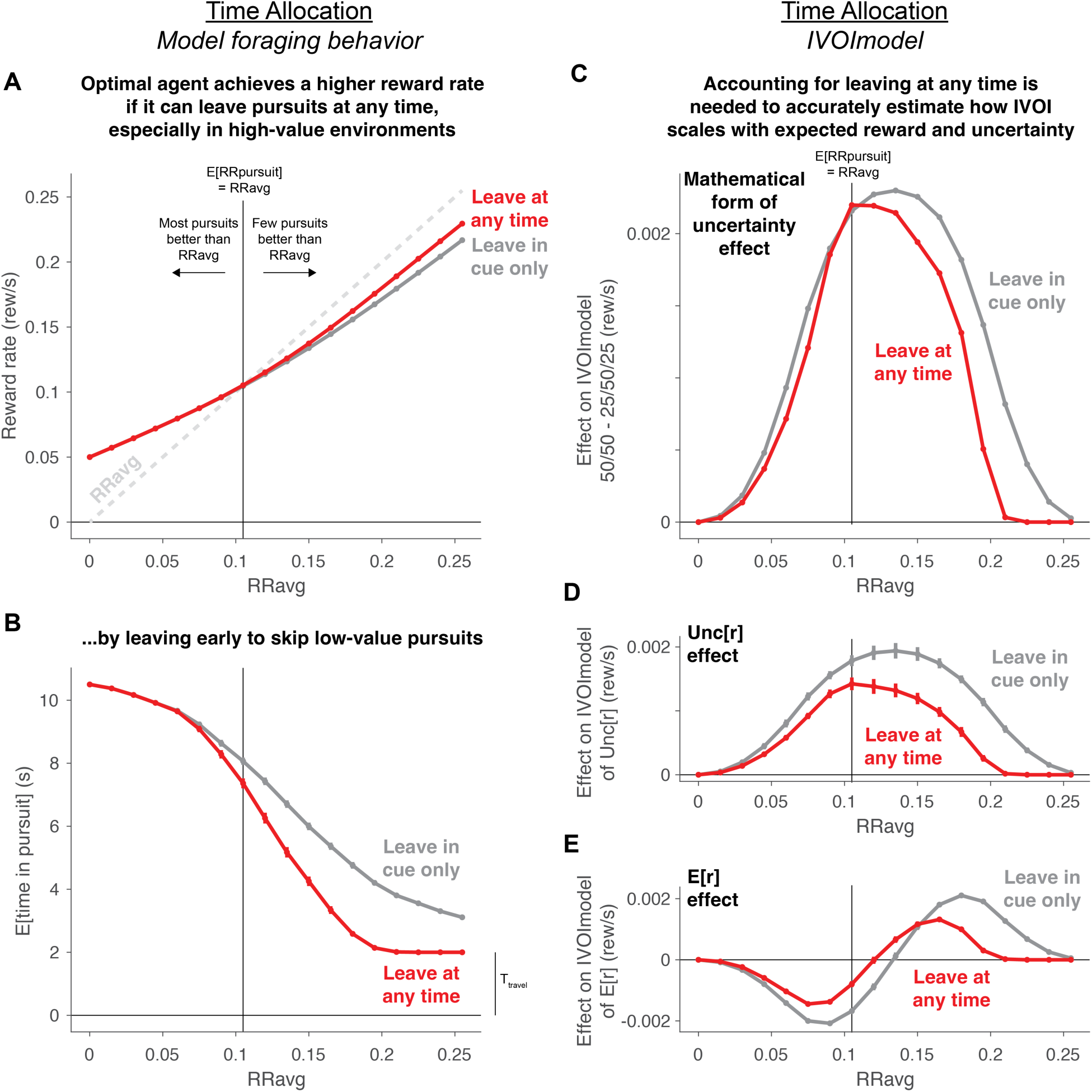
TA theory makes distinct predictions if agents are allowed to leave at any time versus only after observing cues. Here we compare optimal agents in the formulation we propose and use in this paper, where the agent is allowed to leave at any time (red curves; Fig 2), versus optimal agents derived using a modified formulation where the agent is only allowed to leave during the cue period, following a prior formulation based on optimal foraging theory that focused on post-cue leaving decisions^37^ (gray curves). These formulations both produce similar basic patterns of information value: IVOImodel grows with uncertainty (Fig S4D) in a manner resembling SD[r] more than other forms of uncertainty like range or entropy (Fig S4C), and can be modulated by E[r] in either a negative or positive manner depending on the background reward rate RRavg (Fig S4E). However, allowing agents to leave at any time, as they can during most pursuits of reward in natural environments, allows them to obtain a higher reward rate (Fig S4A) due to different leaving behavior (Fig S4B) resulting in different modulations of information value (Fig S4C-E), especially in environments with high background reward rate (data points to the right of the vertical black line in each panel). **(A)** Reward rate of agents that can leave at any time (red) or only during the cue period (gray) as a function of background reward rate (RRavg, x-axis). Vertical black line indicates when RRavg is equal to the mean available reward divided by the time required to pursue it (E[RRpursuit] = RRavg). In each environment, agents are presented with the same set of offers as in Fig 4. Both agents get similar reward rates in environments where RRavg is low, but the agent that can leave at any time gets a higher reward rate in environments where RRavg is high (red > gray). **(B)** Agents that can leave at any time achieve a higher reward rate by leaving early to skip low-value pursuits, especially in high-value environments. Shown is the expected time spent in pursuit (y-axis, including travel time after leaving (T_travel_)) as a function of RRavg (x-axis). Both agents spend less time in pursuits in rich environments with high RRavg. However, this is much more strongly the case for the agent that can leave at any time. The agent saves more than a second of additional time by immediately leaving pursuits that have low E[r], without having to wait for the informative cue to indicate the exact reward size. **(C-E)** Accounting for agents leaving at any time is required to accurately estimate how IVOI scales with the mathematical form of uncertainty (C, curve indicates the difference between 50/50 – 25/50/25 effects from Fig 3B), overall level of uncertainty as measured by SD[r] (D, curve indicates the Unc[r] effect in Fig 3A), and mean offered reward measured by E[r] (E, curve indicates the E[r] effect in Fig 3A), especially in environments with high RRavg (x-axis). **(C)** Both agents scale up information value with a mathematical form of uncertainty that more closely resembles SD[r] than range or entropy, indicated by higher information value for 50/50 offers than 25/50/25 offers (both curves have all y > 0, rather than y = 0 or y < 0). However, the agents only agree on the specific IVOI (red = gray) in an environment where RRavg is equal to the mean pursued reward (vertical black line) – in other words, an environment where the only available reward occurs through this type of pursuit. In other environments, especially rich environments with high RRavg, the agent that can leave at any time assigns a substantially lower IVOI (red < gray). This is because the agent achieves a higher reward rate by sometimes leaving immediately after seeing an offer’s E[r], without waiting for the cue to get information about the exact reward size, and thus it places lower IVOI on the cue. **(D)** Both agents scale up information value with uncertainty as measured by SD[r] (both curves have all y > 0). However, the agent that can leave at any time has uniformly lower IVOI, especially in rich environments with high RRavg, for the same reason discussed above (red < gray). **(E)** Both agents scale down information value with E[r] in low-value environments (low RRavg, left) and scale up information value with E[r] in high-value environments (high RRavg, right). However, the agent that can leave at any time has a shifted curve with lower amplitude modulations in both positive and negative directions, especially in rich environments with high RRavg.

**Figure S5.**
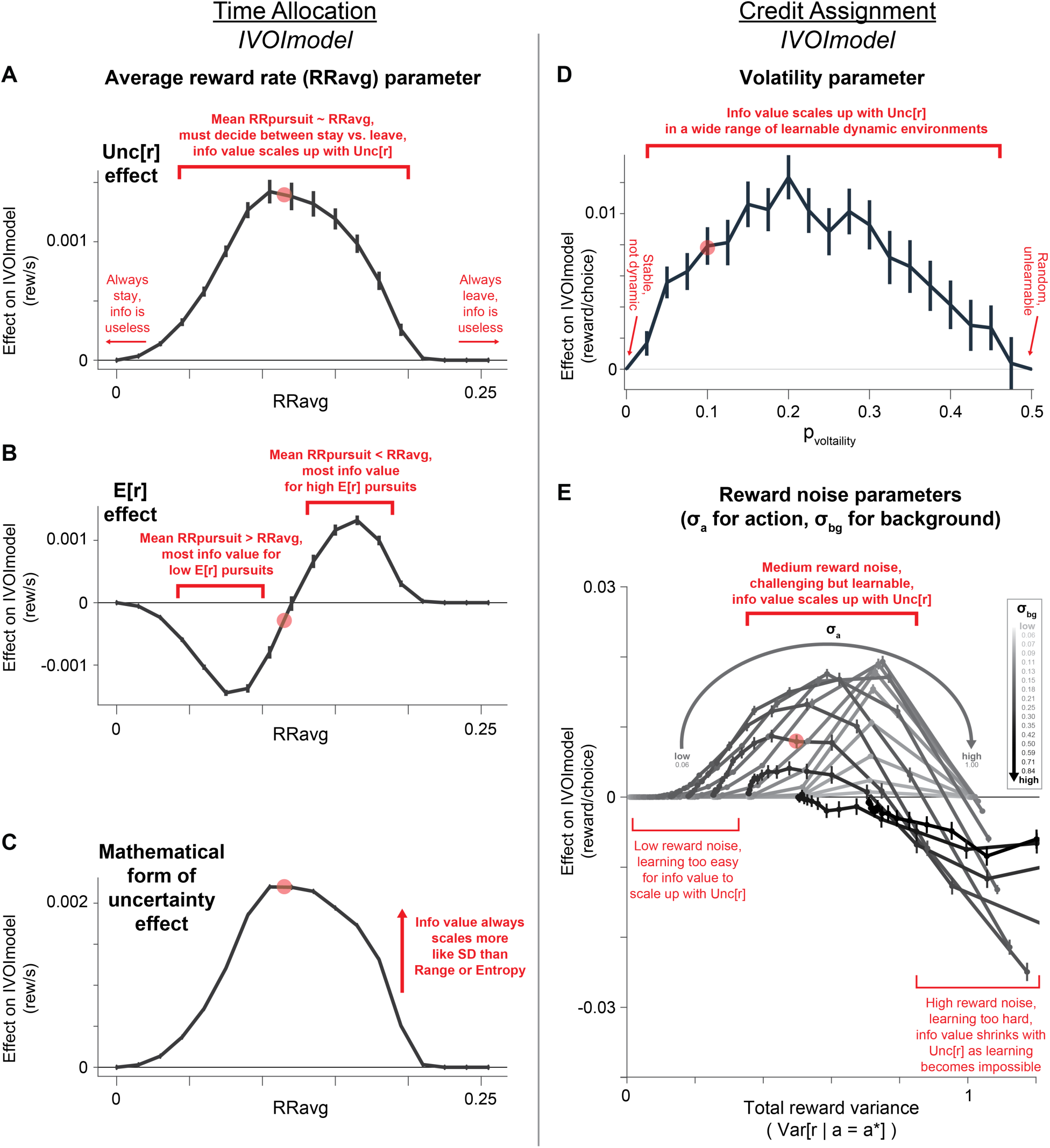
TA and CA model predictions are robust to a wide range of parameters. **(A) TA model consistently scales up IVOImodel with Unc[r].** Each data point on the curve is the same as the Unc[r] effect in Fig 3A, but computed separately for many different settings of RRavg (x-axis). Red circle indicates the setting of RRavg used in Fig 3. IVOImodel consistently scales up with Unc[r] in a wide regime where the mean reward rate of pursuit is roughly similar to the environment’s RRavg (central red bar). This relationship only fails in extreme regimes where RRavg is very low or very high compared to the average reward rate of pursuits (left and right sides of plot, red arrows). This is because information becomes useless for stay/leave decisions in a very poor environment where RRavg is very low (left). The agent will almost always stay in a pursuit, even if the informative cue gives ‘bad news’, because even a bad pursuit still gives a higher reward rate than RRavg. Similarly, information becomes useless in a very rich environment where RRavg is very high (right), because the agent will almost always leave the pursuit even if the informative cue gives ‘good news’. **(B) TA model scales more consistently with Unc[r] than E[r].** Same as A, but showing the E[r] effect. The Unc[r] effect is generally larger than the E[r] effect for most settings of RRavg. Also, unlike the Unc[r] effect which is uniformly positive for all RRavg, the E[r] effect can be either negative, zero, or positive depending on the exact setting of RRavg. This is because IVOImodel is highest when information helps the agent decide whether to stay or leave, which means it is most helpful when there is a close match between RRavg and the E[r] of the reward being pursued. Thus, in poor environments with low RRavg, information is useless for most pursuits because even ‘bad news’ is usually better than abandoning the pursuit. So information is only useful for the subset of pursuits with relatively low E[r] (left red bar, “most info value for low E[r] pursuits”). Conversely, in rich environments with high RRavg, information is useless for most pursuits because even ‘good news’ is usually worse than abandoning the pursuit. So information is only useful for the subset of pursuits with relatively high E[r] (right red bar, “most info value for high E[r] pursuits”). Finally, in moderate environments where information is generally highly useful (Fig S5A), IVOImodel has little net effect of E[r] (middle, y ∼ 0). **(C) TA model scales up IVOImodel with a consistent mathematical form of uncertainty.** Each data point on the curve is the same as the difference between the 50/50 and 25/50/25 effects from Fig 3B, but computed separately for different settings of RRavg (x-axis). Thus, IVOImodel scales more like an SD-like, range-like, or entropy-like form of uncertainty if data points have y > 0, y = 0, or y < 0. All data points have y > 0 (except for extreme settings where information is useless and hence IVOImodel is zero; left and right sides of plot), indicating that information value consistently scales with an SD-like form of uncertainty. **(D) CA model scales up IVOImodel consistently for a wide range of environmental volatilities.** Same as C, but for the CA model and computed separately for different settings of p_volatility_ (x-axis). IVOImodel scales up positively in an SD-like manner for all settings that are learnable dynamic environments, with a broad peak including a wide range of high and low volatility levels (central red bar). The only situations where IVOImodel does not scale up are extreme regimes that are not learnable dynamic environments. This occurs when volatility is at the minimum possible setting, near-zero, so the environment is not dynamic (left side, red arrow). This also occurs when volatility is at the maximal possible setting, near 0.5, so the environment is totally random and not learnable, such that the good and bad actions are completely randomized before each choice (right side, red arrow). **(E) CA model scales up IVOImodel with reward noise, arising both from actions and background events, for a wide range of noise levels.** Same as D, but computed separately for different settings of the reward noise arising from the action (σ_a_) and arising from background events (σ_bg_). Data is plotted as a function of the total empirical reward variance arising from both sources (x-axis, Var[*r* | *a* = *a*\*]). σ_a_ and σ_bg_ were each varied independently testing all possible settings in a wide range spaced equally on a log axis. Each curve represents a different setting of σ_bg_, with the curve’s shading indicating the precise setting (indicated by the legend; ranging from σ_bg_ = 0.06 in light gray to σ_bg_ = 0.84 in black). Each data point on the curve represents a different setting of σ_a_ (indicated by gray arrow; ranging from σ_a_ = 0.06 on the left to σ_a_ = 1 on the right). We find that IVOImodel generally scales up with uncertainty in a similar manner for a wide range of intermediate reward noise levels, including reward noise arising from both sources (central red bar, “Medium reward noise”). While the exact IVOImodel depends on the exact settings of σ_a_ and σ_bg_ (e.g. σ_bg_ has a slightly larger effect in this regime than σ_a_), but these curves generally have a broad positive peak in a similar region of intermediate reward noise. Thus, in this region, the agent places higher value on information when either actions or background events have higher reward variance. That is when information is most necessary for the agent, in to combat the reward variance and continue to learn efficiently. Of course, as with volatility, this relationship is not absolute. Extreme settings of either noise parameter can eliminate information value, causing IVOImodel effects to become near-zero or even negative (red bars on left and right sides of plot). In particular, if reward noise is very low, then IVOImodel has close to zero scaling with uncertainty (left side, “Low reward noise”, y ∼ 0). This is because learning becomes so easy that information is useless, since the agent can learn almost perfectly even without information. On the other hand, if reward noise is very high, then IVOImodel has zero or even negative scaling with uncertainty (right side, “High reward noise”, y <= 0). This occurs especially when σ_bg_ is high, for which the curves are entirely negative (blackest curves). This is because learning becomes so hard that information is little help to the agent. Adding more reward variance just makes learning increasingly impossible, and hence makes information even less helpful, thus having a negative effect on IVOImodel.

**Figure S6.**
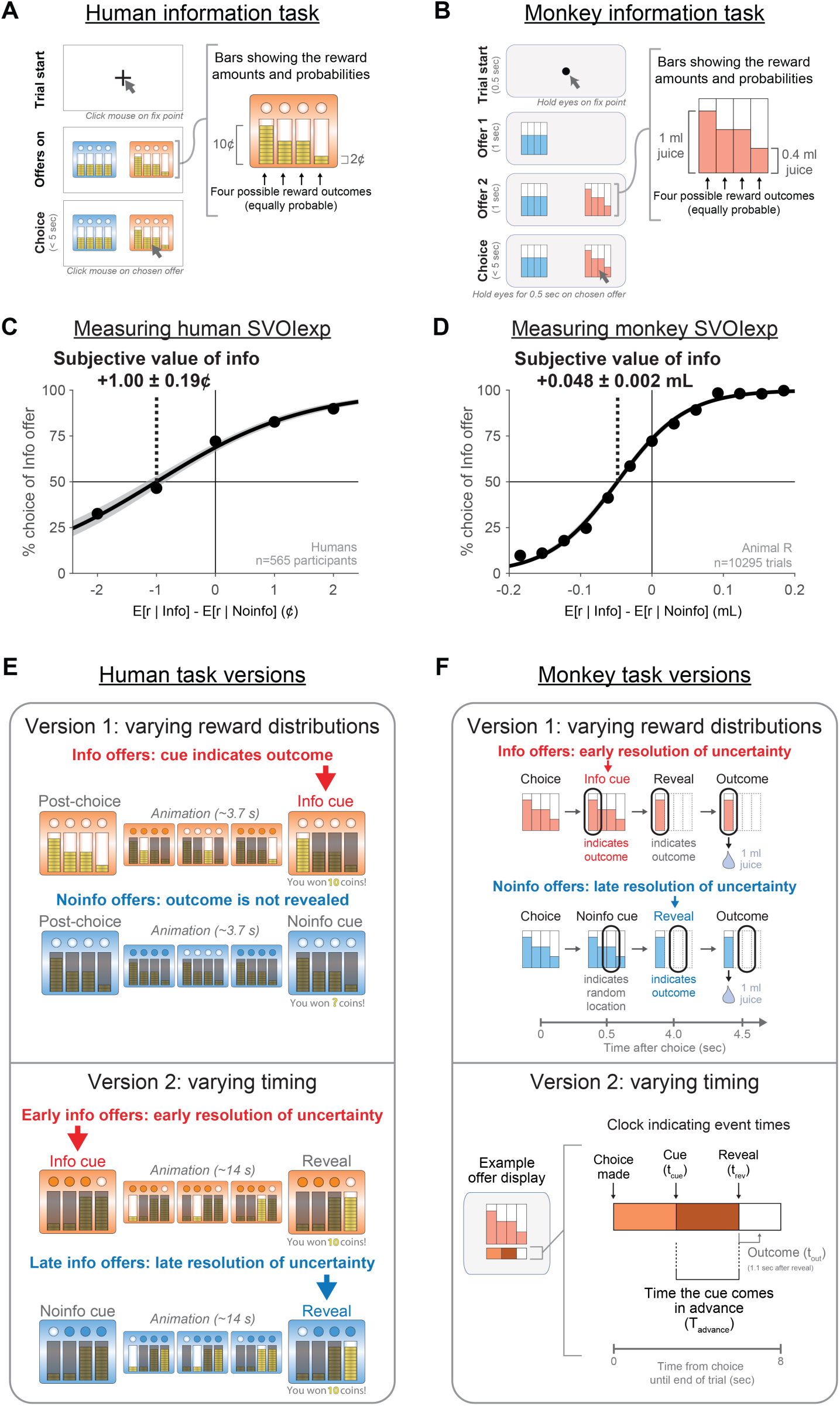
Experimental designs of analogous information seeking tasks for humans and monkeys. **(A) Multi-attribute information choice task for humans.** Same task design and data as^8^. Humans used mouse clicks to choose between a pair of offers that each explicitly showed their probability distribution of reward outcomes, in the form of four possible equally probable reward outcomes whose sizes were indicated by stacks of coins. Offers with different colored textures were either Info offers that provided informative cues or Noinfo offers that did not. **(B) Multi-attribute information choice task for monkeys.** Same task design and data as^8^. Monkeys used eye movements to choose between a pair of offers that each explicitly showed their probability distribution of reward outcomes, in the form of four possible equally probable reward outcomes whose sizes were indicated by colored, textured bars. Offers with different colored textures were either Info offers that provided informative cues or Noinfo offers that did not. Note that the diagrams here use simple red and blue colors for illustrative purposes, but the bars used in experiments were shown using textures derived from natural scenes. **(C,D) Measuring SVOIexp in humans and monkeys.** We measured the subjective value of information in the standard economic sense of how much reward individuals willingly paid to gain the information. This can be visualized with a simple psychophysics plot of the percent choice of Info as a function of the difference in E[r] between the Info and Noinfo offers (using only the subset of trials where one offer was Info and the other was Noinfo). The mean data are shown from the human population (n=565, left) and from an example monkey (animal R, right), with black curves indicating logistic fits to the data. These plots in both species indicate a strong preference for Info. For instance, they both had > 70% choice of Info when the two offers had equal E[r] (y > 70% when x = 0). To measure SVOIexp, we used the logistic fits to the data (black curves) to estimate the *indifference point* – the difference in E[r] favoring the Noinfo offer that exactly counterbalanced the individual’s inherent preference for Info, and caused them to be indifferent between the two offers (black dashed line). In these examples, this measure of SVOIexp was 1.00 ± 0.19 cents in humans and 0.048 ± 0.002 mL juice in the monkey (mean ± bootstrap SE). This illustrates how we can measure SVOIexp in these experiments allowing individuals to trade reward for information. Of course, in these tasks offers could vary in many different factors at once, not just a single factor E[r]. Therefore, our analysis in this work and our previous studies generally has to estimate SVOIexp using more sophisticated methods. Most commonly, we fit choice data using descriptive models (GLMs) that can simultaneously incorporate the effects of many different factors and their interactions on subjective value (Methods). **(E,F) Human and monkey tasks varying reward distributions (version 1) and event timing (version 2).** **(E)** *Top:* Human task version 1 focused on varying reward distributions and their effects on SVOIexp. An offer’s color indicated its informativeness. Info offers provided participants with a short animation followed by an informative cue indicating which of the four possible outcomes would be delivered, including a text cue (e.g. “You won 10 coins!”). Noinfo offers provided participants with no such stimuli and a non-informative text cue (e.g. “You woncoins!”), thus leaving the reward outcome uncertain. Participants were instructed that regardless of whether they saw informative or non-informative cues, the reward outcome was still deposited into their winnings at the end of the trial. Thus, Info was non-instrumental: it only allowed participants to know the outcome they would receive, but did not allow them to control that outcome. *Bottom:* Human task version 2 focused on varying timing and its effect on SVOIexp. In this task version, Early Info offers provided a cue indicating the upcoming reward outcome immediately after the choice, while Late Info offers provided a non-informative cue that indicated a random stack of coins and hence left the outcome uncertain. Then, both offers played identical animations followed by identical final reveals that indicated the reward outcome. Thus, both offer types eventually indicated the true outcome, but Early Info offers indicated the outcome much earlier in advance than Late Info offers. **(F)** *Top:* Monkey task verson 1 focused on varying reward distributions and their effects on SVOIexp. Info offers provided an informative cue that indicated which of the four possible outcomes would be delivered, while Noinfo offers provided a non-informative cue that indicated a random bar and hence left the outcome uncertain. Note that unlike the human task, the monkeys worked for juice and hence had to have the reward physically delivered on each trial. Thus, for both Info and Noinfo offers, we provided an additional ‘reveal’ stimulus that always indicated the outcome 0.5 sec before reward delivery, to ensure that animals had always had appropriate time to physically prepare to receive the reward regardless of whether they chose Info or Noinfo ^6,8^. *Bottom:* Monkey task version 2 focused on varying timing and its effect on SVOIexp. In this task version, each offer was augmented with an additional horizontal bar at the bottom of the offer stimulus (left), that functioned as a “clock” indicating event timings. The horizontal length of the bar represented the time duration from choice to the end of the trial. The bar was split into three segments each covered with a unique colored texture. The first segment indicated the time from choice to cue (t_cue_); the second indicated time from cue to reveal (t_reveal_); and the third indicated time from reveal until the end of the trial. The reveal always occurred 1.1 s in advance of outcome delivery (t_out_). After choice, an animated ‘hand of the clock’ continuously showed the passage of time, and successively indicated when the cue and reveal events were triggered when it touched the boundaries between bar segments. Offers had varying t_cue_ and t_reveal_, allowing us to determine how SVOIexp varied with the timing of the cue, the reveal, and their difference, the time the information came in advance (T_advance_; Methods).

**Figure S7.**
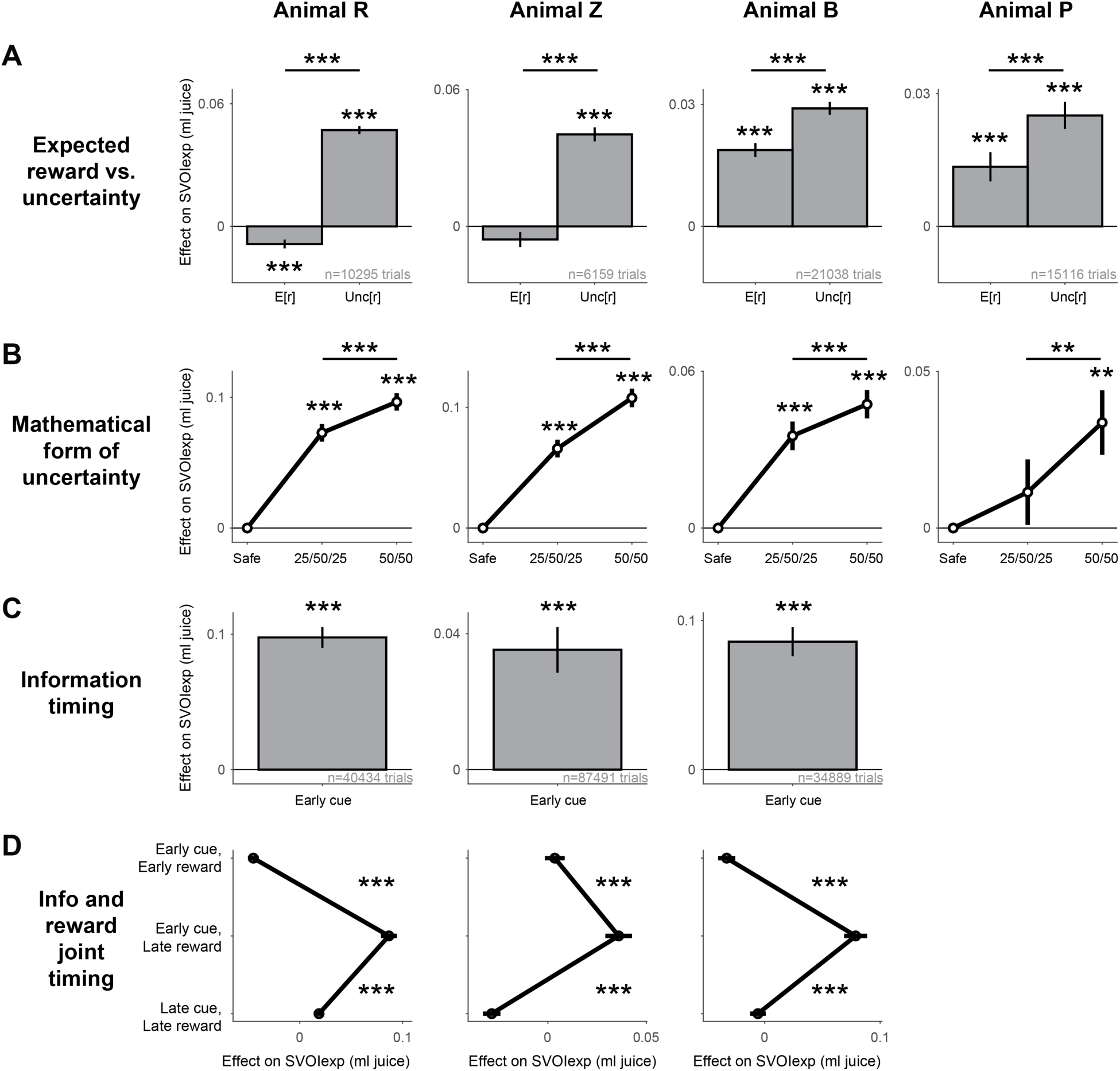
All monkeys scaled SVOIexp in consistent manners with multiple factors including expected reward, uncertainty, and cue and reward timing. **(A,B)** Same results as Fig 3A,B separately for each of the n=4 animals that were tested with version 1 of the monkey task (manipulating reward distributions with fixed timing), as reported previously ^8^. All showed the same qualitative pattern of results. SVOIexp is significantly positively related to Unc[r] (A, Unc[r] weight > 0, all p < 0.001), significantly more positively related to Unc[r] than E[r] (A, Unc[r] weight > E[r] weight, all p < 0.001). SVOIexp scales up with uncertainty in a manner significantly more similar to SD[r] than to alternate measures like range or entropy, indicated by higher SVOIexp for 50/50 reward distributions than 25/50/25 distributions (B, all p < 0.01). Also, different animals scaled SVOIexp with E[r] in a manner that was either negative, non-significant, or positive (animal R, negative; animal Z, non-significant; animals B and P, positive). This resembles the predictions of the TA theory, which predicts that an agent may have negative, zero, or positive effects of E[r] on the agent’s estimate of the environment’s RRavg (Fig S4, Fig S5). **(C,D)** Same results as Fig 3C,D, separately for each of the n=3 animals that were tested with version 2 of the monkey task (manipulating reward distributions and cue and reward timing), as reported previously ^8^. All showed the same qualitative pattern of results. SVOIexp was significantly positively related to cue timing (C, higher for early cues than late cues, all p < 0.001) and significantly higher when the cue came early in advance of the outcome (D, higher for “early cue, late reward” than “early cue, early reward” and “late cue, late reward”, all p < 0.001).

**Figure S8.**
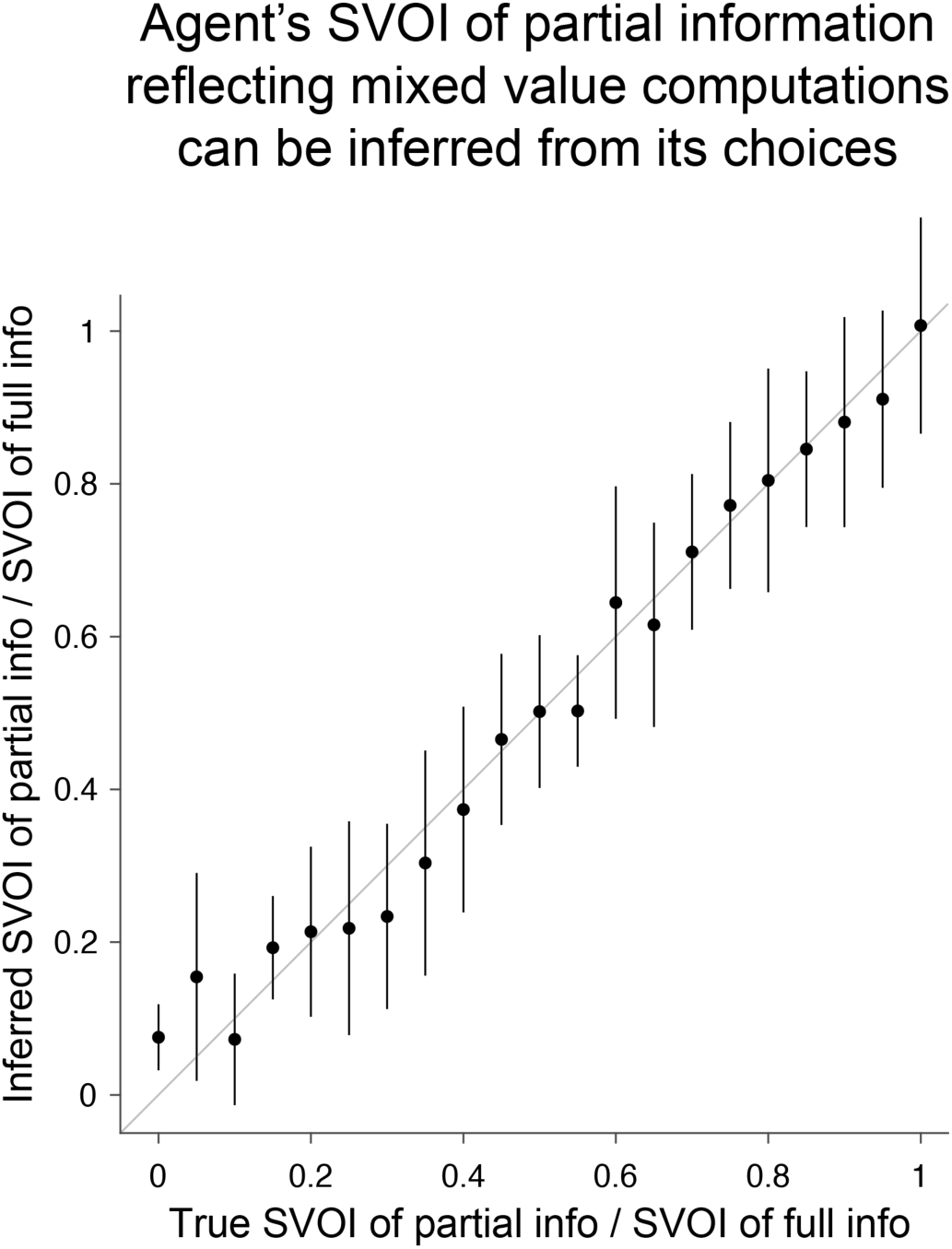
An agent’s SVOI of partial information, reflecting its mixed value computations, can be inferred from its choices. Shown is the relationship between true (x-axis) and inferred (y-axis) SVOI of partial information, expressed as a fraction of SVOI of full information. Thus, this quantity is 0 if the agent is unwilling to pay for partial information, and 1 if the agent is willing to pay just as much for partial information as full information. SVOIs were inferred based on the estimated cost the agent was willing to pay for partial and full information based on its choice data. Error bars are ±SD across all n=11 environments for simulated agents with true partial/full SVOI fractions ranging from 0 to 1 in increments of 0.05. In all cases, the inferred quantity closely matches the true quantity (x ≈ y).

**Table S1.**
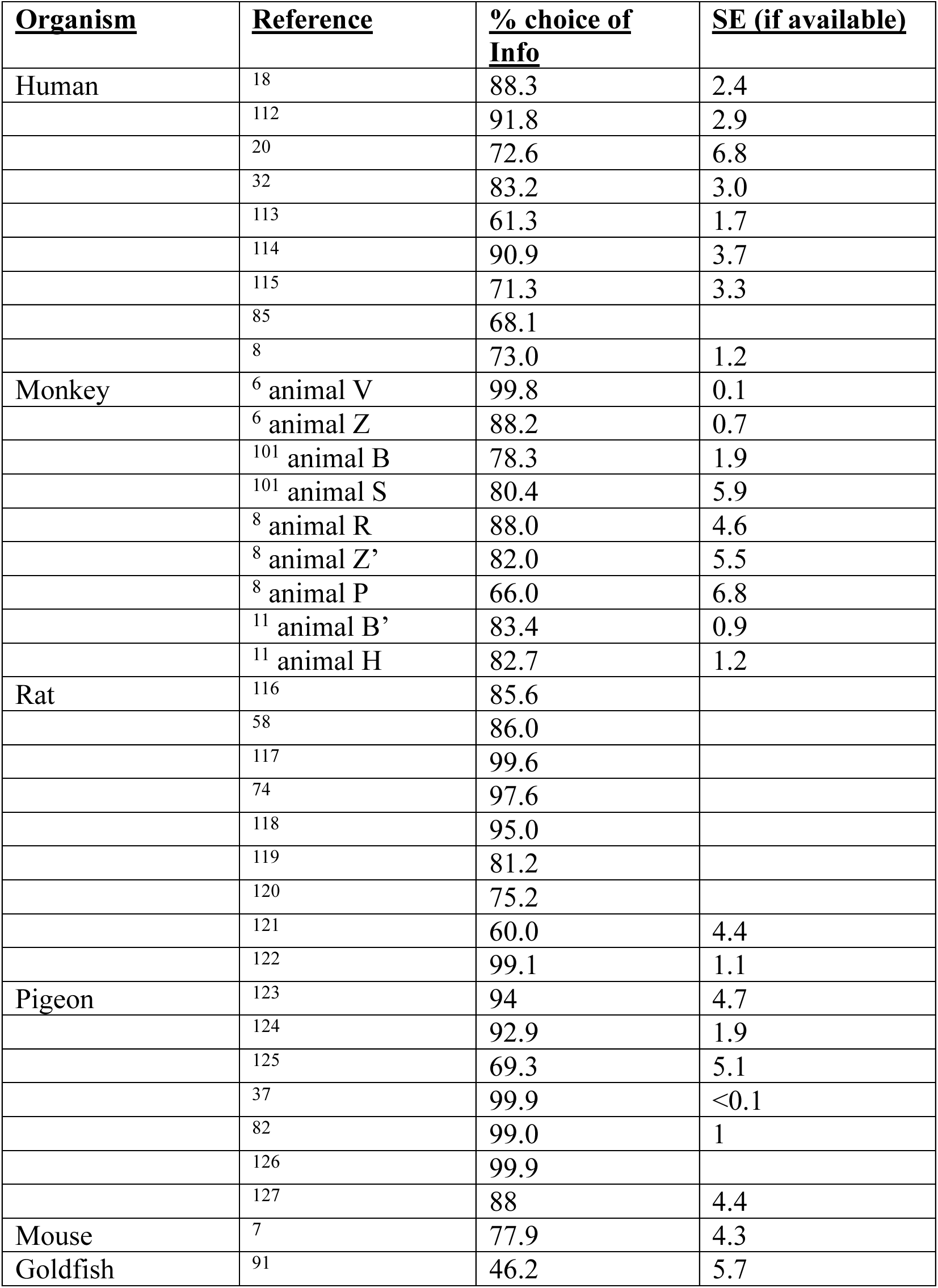
References used for plot of percent choice of information across species. As we discuss in the main text, a rigorous comparison of information preferences across species requires use of experimental tasks that are as analogous as possible and measurement of subjective values rather than only percent choice. However, such data is available in few cases^8,89^. In the absence of such data, it is still possible to use percent choice to illustrate whether individuals of a given species tend to prefer Info, to prefer Noinfo, or to be indifferent between the two, without making strong claims about the relative degree of those preferences across species. Hence, this is what the corresponding plot here is used to illustrate (Fig 1B). We used a set of studies that all meet the following criteria to ensure they are as comparable as possible given these limitations. (1) Must measure percentage choice between Info versus Noinfo options analogous to those in Fig 1A. (2) Those options must only have outcomes that are appetitive (monetary for humans, typically food or water for animals) or neutral, not aversive. (3) Must have uncertain rewards – that is, p(reward) > 0% and < 100%; if multiple probabilities were tested, we use data from condition closest to p(reward) = 50%. (4) Must have both good and bad possible outcomes indicated by different informative cues, which could be in the form of different reward sizes or time delays or effort requirements to obtain the reward; we use data from the condition with the greatest difference between good and bad outcomes (e.g. biggest reward size difference). (5) Must have delayed rewards; we use data from the greatest reward delay tested. (6) Must have informative cues that are valid; we use data from the highest validity tested. (7) Must have Info and Noinfo cues with matched stimulus properties; for example, if there are two possible Info cues, then there must be two matched Noinfo cues with corresponding probabilities and similar physical characteristics. (8) Info and Noinfo options must have the same reward distribution; except in pigeons where the great majority of choice experiments use asymmetric rewards based on an early task design^9^. (9) If data is presented from multiple days of training, use data from the last day of training. Choice percentages and standard errors were measured either from reported results in text format or estimated algorithmically from data plots in figures; or from primary datasets when possible. Data in each row for monkeys is from an individual animal and is labeled with that animal’s descriptor used in the original study, and each row is a different animal (rows with the same descriptor correspond to different animals from different studies that happened to be given the same descriptor). Note that these criteria are not meant to encompass all types of information seeking; other types of studies examine other aspects of information seeking (e.g. different reward distributions for Info and Noinfo ^128^, responses other than direct choices between Info and Noinfo such as response rates on variable-interval schedules^2^, properties of saccades ^129^ or curiosity ratings ^19^, information about counterfactual outcomes rather than actual outcomes ^78^, etc.).

| <u>Organism</u> | <u>Reference</u> | <u>% choice of Info</u> | <u>SE (if available)</u> |
| --- | --- | --- | --- |
| Human | 18 | 88.3 | 2.4 |
|  | 112 | 91.8 | 2.9 |
|  | 20 | 72.6 | 6.8 |
|  | 32 | 83.2 | 3.0 |
|  | 113 | 61.3 | 1.7 |
|  | 114 | 90.9 | 3.7 |
|  | 115 | 71.3 | 3.3 |
|  | 85 | 68.1 |  |
|  | 8 | 73.0 | 1.2 |
| Monkey | <sup>6</sup> animal V | 99.8 | 0.1 |
|  | <sup>6</sup> animal Z | 88.2 | 0.7 |
|  | <sup>101</sup> animal B | 78.3 | 1.9 |
|  | <sup>101</sup> animal S | 80.4 | 5.9 |
|  | <sup>8</sup> animal R | 88.0 | 4.6 |
|  | <sup>8</sup> animal Z' | 82.0 | 5.5 |
|  | <sup>8</sup> animal P | 66.0 | 6.8 |
|  | <sup>11</sup> animal B' | 83.4 | 0.9 |
|  | <sup>11</sup> animal H | 82.7 | 1.2 |
| Rat | 116 | 85.6 |  |
|  | 58 | 86.0 |  |
|  | 117 | 99.6 |  |
|  | 74 | 97.6 |  |
|  | 118 | 95.0 |  |
|  | 119 | 81.2 |  |
|  | 120 | 75.2 |  |
|  | 121 | 60.0 | 4.4 |
|  | 122 | 99.1 | 1.1 |
| Pigeon | 123 | 94 | 4.7 |
|  | 124 | 92.9 | 1.9 |
|  | 125 | 69.3 | 5.1 |
|  | 37 | 99.9 | <0.1 |
|  | 82 | 99.0 | 1 |
|  | 126 | 99.9 |  |
|  | 127 | 88 | 4.4 |
| Mouse | 7 | 77.9 | 4.3 |
| Goldfish | 91 | 46.2 | 5.7 |

**Table S2.** GLM regressors for each dataset, task version, and analysis. Continuous factors are defined in text (E[r], SD[r]). Binary indicator variables are defined as follows: Left (1 for offers on the right side of the screen and 0 otherwise), Right (1 for offers on the right side and 0 otherwise), Second (1 for the offer that was presented second in the trial, 0 for the offer that was presented first), Info (for Info term: +0.5 for Info offers and -0.5 otherwise; for interactions: 1 for Info and 0 otherwise), 25/50/25 and 50/50 (1 for the specified offer type, 0 otherwise), EE (1 for offers with early cue and early outcome, 0 otherwise), EL (1 for offers with early cue and late outcome, 0 otherwise), LL (1 for offers with late cue and late outcome, 0 otherwise). Cue (or outcome) times were classified as early or late if they occurred in the 1^st^ or 3^rd^ tercile of cue (or outcome) times, Constant (1 always).

| Dataset | Task version | Analysis | Regressors |
| --- | --- | --- | --- |
| Human | 1 | E[r] vs Unc[r] | E[r], SD[r], Info, Info x E[r], Info x SD[r], Right |
| Human | 1 | SD vs Range vs Entropy | E[r], 25/50/25, 50/50, Info, Info x E[r], Info x 25/50/25, Info x 50/50, Right |
| Human | 2 | Early vs Late cue | E[r], EarlyInfo, Right, Constant |
| Monkey | 1 | E[r] vs Unc[r] | E[r], SD[r], Info, Info x E[r], Info x SD[r], Left, Right, Second |
| Monkey | 1 | SD vs Range vs Entropy | E[r], 25/50/25, 50/50, Info, Info x E[r], Info x 25/50/25, Info x 50/50, Left, Right, Second |
| Monkey | 2 | Early vs Late cue | E[r], SD[r], $t_{outcome}$, E[r] x SD[r], Info, Info x SD[r], Info x (EE – LL), Info x LL, Left, Right, Second |
| Monkey | 2 | Info and Reward timing | E[r], SD[r], $t_{outcome}$, E[r] x SD[r], Info, Info x SD[r], Info x EE, Info x EL, Info x LL, Left, Right, Second |
| Model | 1 | E[r] and Unc[r] | E[r], SD[r], E[r] x SD[r], Constant |
| Model | 1 | SD vs Range vs Entropy | E[r], 25/50/25, 50/50, E[r] x 25/50/25, E[r] x 50/50, Constant |
| Model | 2 | Early vs Late cue | E[r], SD[r], E[r] x SD[r], EL-LL, LL, Constant |
| Model | 2 | Info and Reward timing | E[r], SD[r], E[r] x SD[r], EE, EL, LL, Constant |

